# Mitochondrial Na^+^ controls oxidative phosphorylation and hypoxic redox signalling

**DOI:** 10.1101/385690

**Authors:** Pablo Hernansanz-Agustín, Carmen Choya-Foces, Susana Carregal-Romero, Elena Ramos, Tamara Oliva, Tamara Villa-Piña, Laura Moreno, Alicia Izquierdo-Álvarez, J. Daniel Cabrera-García, Ana Cortés, Ana Victoria Lechuga-Vieco, Pooja Jadiya, Elisa Navarro, Esther Parada, Alejandra Palomino-Antolín, Daniel Tello, Rebeca Acín-Pérez, Juan Carlos Rodríguez-Aguilera, Plácido Navas, Ángel Cogolludo, Iván López-Montero, Álvaro Martínez-del-Pozo, Javier Egea, Manuela G. López, John W. Elrod, Jesús Ruiz-Cabello, Anna Bogdanova, José Antonio Enríquez, Antonio Martínez-Ruiz

## Abstract

All metazoans depend on O_2_ delivery and consumption by the mitochondrial oxidative phosphorylation (OXPHOS) system to produce energy. A decrease in O_2_ availability (hypoxia) leads to profound metabolic rewiring. In addition, OXPHOS uses O_2_ to produce reactive oxygen species (ROS) that can drive cell adaptations through redox signalling, but also trigger cell damage^1–4^, and both phenomena occur in hypoxia^4–8^. However, the precise mechanism by which acute hypoxia triggers mitochondrial ROS production is still unknown. Ca^2+^ is one of the best known examples of an ion acting as a second messenger^9^, yet the role ascribed to Na^+^ is to serve as a mere mediator of membrane potential and collaborating in ion transport^10^. Here we show that Na^+^ acts as a second messenger regulating OXPHOS function and ROS production by modulating fluidity of the inner mitochondrial membrane (IMM). We found that a conformational shift in mitochondrial complex I during acute hypoxia^11^ drives the acidification of the matrix and solubilization of calcium phosphate precipitates. The concomitant increase in matrix free-Ca^2+^ activates the mitochondrial Na^+^/Ca^2+^ exchanger (NCLX), which imports Na^+^ into the matrix. Na^+^ interacts with phospholipids reducing IMM fluidity and mobility of free ubiquinone between complex II and complex III, but not inside supercomplexes. As a consequence, superoxide is produced at complex III, generating a redox signal. Inhibition of mitochondrial Na^+^ import through NCLX is sufficient to block this pathway, preventing adaptation to hypoxia. These results reveal that Na^+^ import into the mitochondrial matrix controls OXPHOS function and redox signalling through an unexpected interaction with phospholipids, with profound consequences in cellular metabolism.

---

Cells and tissues produce a superoxide burst as an essential feature of several adaptive responses, including hypoxia^7,11^. Given the importance of NCLX in ischemia-reperfusion injury^12^ we studied whether this exchanger may have a role in hypoxic redox signalling. Primary bovine aortic endothelial cells (BAECs) and mouse embryonic fibroblasts (MEFs) exposed to acute hypoxia showed an increase in cytosolic Ca^2+^ (Ca^2+^_cyto_) and a decrease in cytosolic Na^+^ (Na^+^_cyto_) that was prevented by NCLX knock-down with siRNAs, genetic deletion, overexpression of a dominant negative form of NCLX (dnNCLX) or pharmacologic inhibition with CGP-37157 (Fig. 1a-e; Extended Data Figs. 1a-j and 2). Importantly this effect was rescued by expression of human NCLX in NCLX KO MEFs (Fig. 1d-e; Extended Data Figs. 1e-g). In agreement with the role of NCLX in mitochondrial Ca^2+^ handling^12^, the transient increase in mitochondrial Ca^2+^ after histamine induction was more persistent upon NCLX down-regulation (Extended Data Fig. 1a-j). Notably, NCLX inhibition did not interfere with mitochondrial membrane potential or respiration (Extended Data Fig 1k-l). Thus, acute hypoxia induces NCLX activation.

**Fig. 1.**
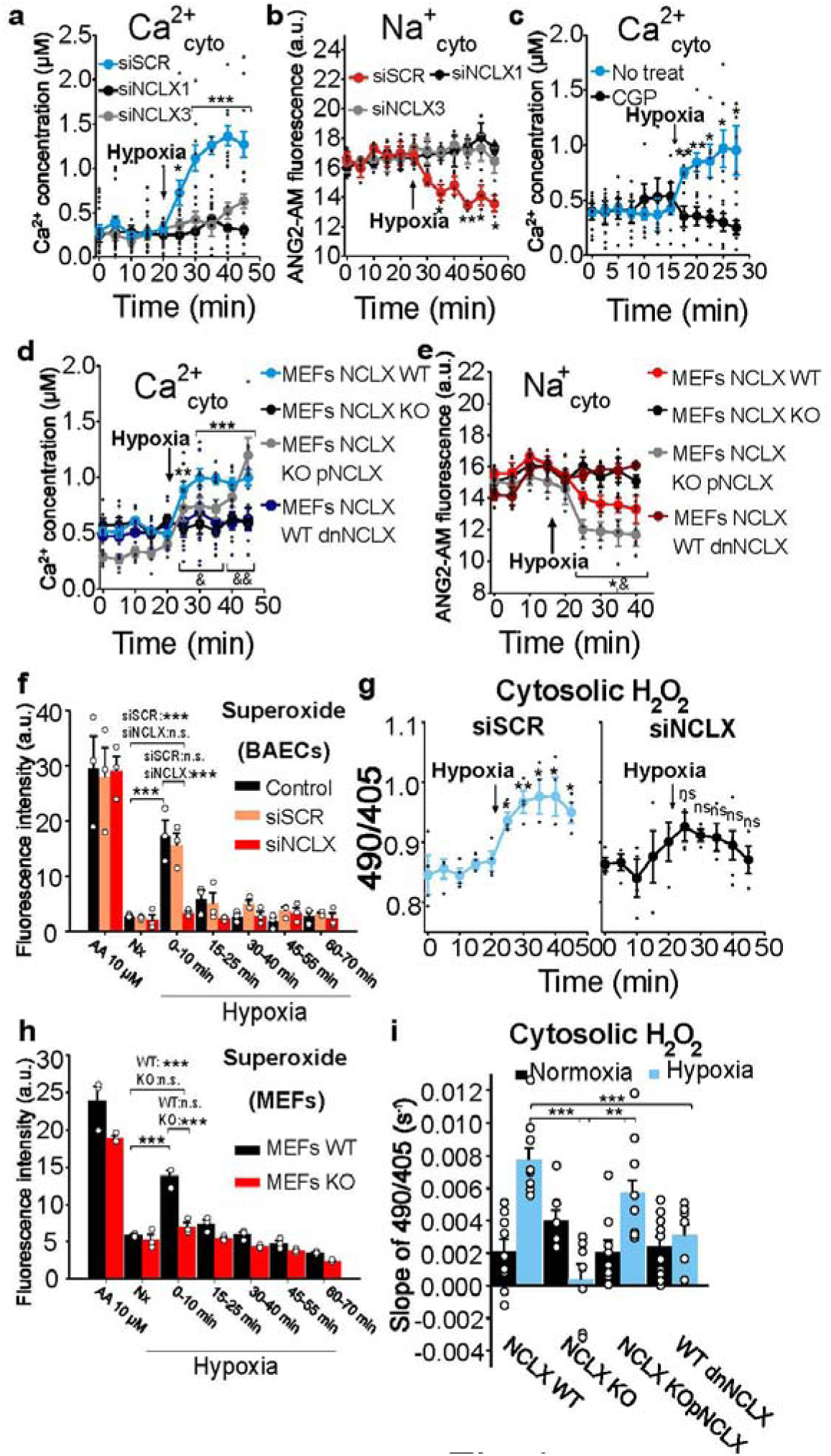
Hypoxia activates Na^+^/Ca^2+^ exchange and enhances ROS production through NCLX. **a-e**, cytosolic Ca^2+^ (Cyto-GEM-GECO; Ca^2+^_cyto_) or Na^+^_cyto_ (Asante Natrium 2-AM; ANG2-AM) measured by live cell confocal microscopy in normoxia and after induction of acute hypoxia (1% O_2_). **a-b**, BAECs transfected with scramble siRNA (siSCR), siNCLX1 or siNCLX3. **c**, Non-treated BAECs (No treat) or treated with CGP-37157. **d-e**, WT, KO, KO+pNCLX or WT+dnNCLX MEFs. **f-i**, ROS production measured by live cell confocal microscopy **(g, i)** or fixed cell fluorescence microscopy **(f, h)** in Nx or acute hypoxia (1% O_2_). **f**, Superoxide detection after incubation with DHE in 10-min time windows in non-treated (Control), siSCR or siNCLX treated cells; AA = antimycin A. **g**, Detection of H_2_O_2_ with CytoHyPer in siSCR- or siNCLX-treated BAECs. **h**, Superoxide detection with DHE in WT or KO immortalized MEFs. **i**, Detection of H_2_O_2_ with CytoHyPer in WT, KO, KO+pNCLX or WT+dnNCLX immortalized MEFs. **a-c**, time-course traces of six independent experiments; **d, e**, time-course traces of four independent experiments; **f, h**, mean intensity of three independent experiments; **g**, time-course traces of four; **i**, slopes of nine independent experiments. One-way ANOVA with Tukey’s test for multiple comparisons (f-h, i) and student’s t-test (last Nx time vs Hp times; a-e, g): * p < 0.05, ** p < 0.01, *** p<0.001. Student’s t-test (WT vs KO): ^&^ p < 0.05, ^&&^ p < 0.01. **f,h**, statistical comparisons shown only for Nx vs 0-10 groups.

**Fig. 2.**
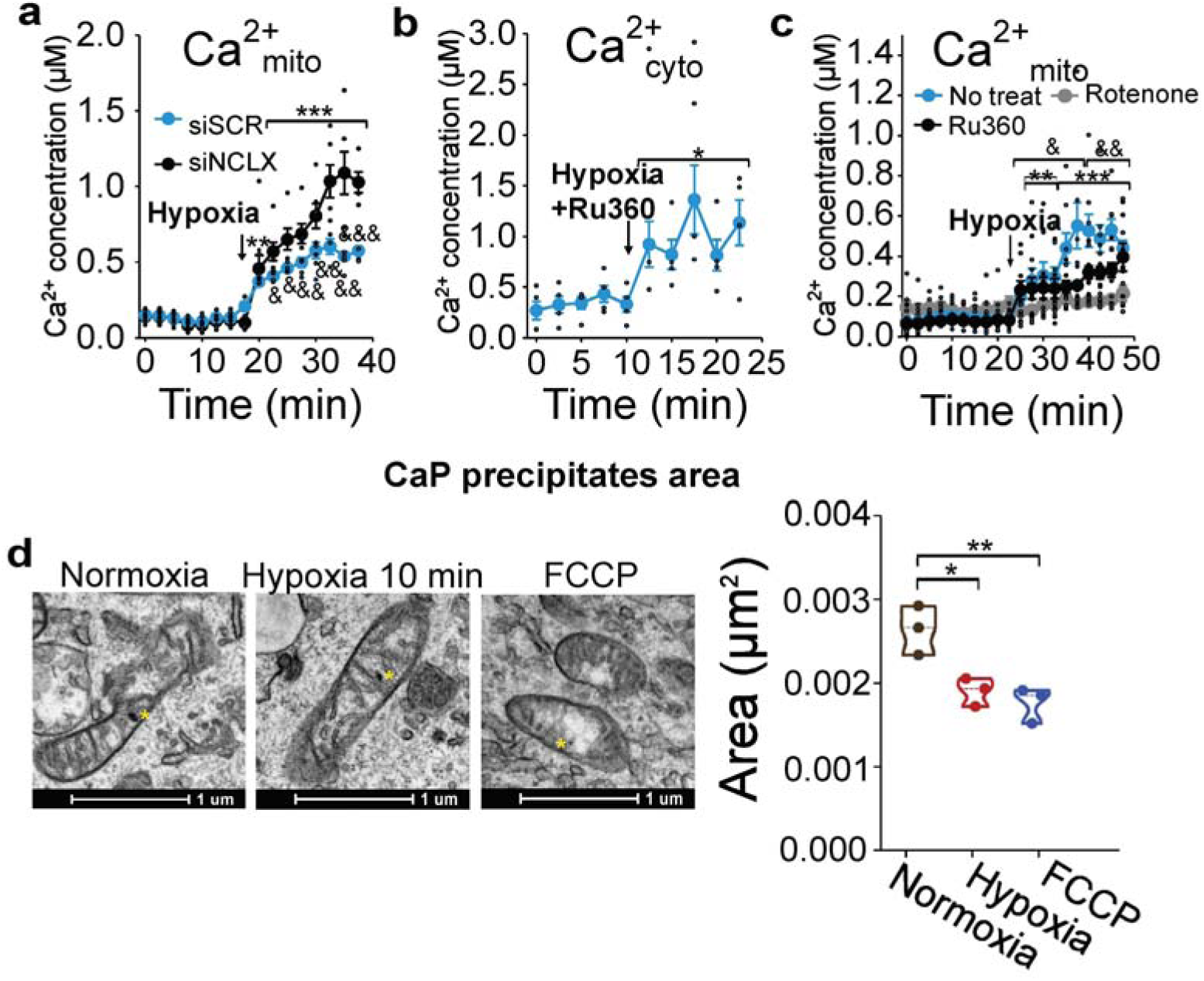
Mitochondrial calcium phosphate precipitates are the source of mitochondrial Ca^2+^ raise and of NCLX activation. **a**, Ca^2+^_mito_ (Cepia 2mt) measured by live cell confocal microscopy in BAECs transfected with siSCR or siNCLX during normoxia (Nx) or acute hypoxia (1% O_2_; trace of four independent experiments). **b**, Ca^2+^_cyto_ measured by live cell confocal microscopy in BAECs subjected to normoxia (Nx) and acute hypoxia (1% O_2_) in the presence of 1 µM Ru360, MCU blocker; trace of four independent experiments). **c**, Ca^2+^_mito_ measured by live cell confocal microscopy in non-treated BAECs or treated with 1 µM Ru360 or 1 µM rotenone during normoxia (Nx) or acute hypoxia (1% O_2_; trace of five or more independent experiments). **d**, Representative TEM images and mean area of mitochondrial calcium phosphate precipitates during normoxia (n=110 precipitates), 10 min hypoxia (1% O_2_; n=145 precipitates) or 30 min treatment with 1 µM FCCP (n=96 precipitates) in three independent experiments. One-way ANOVA with Tukey’s test for multiple comparisons (d) and student’s t-test (last Nx time vs Hp times; a-c): * p<0.05, **p<0.01, *** p<0.001. Student’s t-test (siSCR vs siNCLX): ^&^ p < 0.05, ^&&^ p < 0.01, ^&&&^ p < 0.001.

The hypoxic ROS burst was blocked by NCLX knock-down by siRNA (from now on referred to as siNCLX), deletion, overexpression of dnNCLX, or inhibition with CGP-37157 (Fig.1f-i and Extended Data Fig. 3); and rescued by expression of human NCLX in KO MEFs (Fig. 1i). Therefore, NCLX activity is necessary for mitochondrial ROS production during hypoxia.

**Fig. 3.**
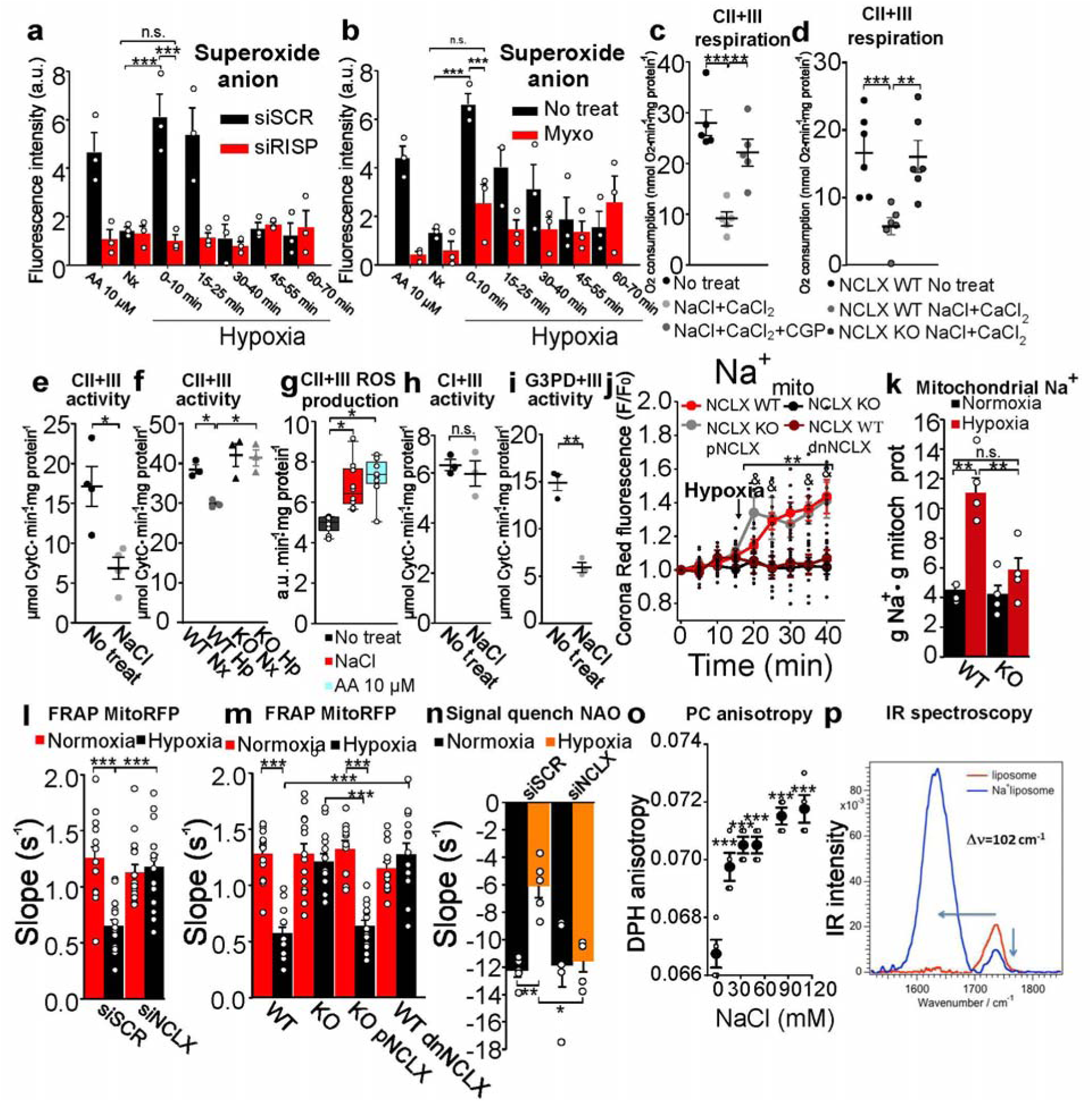
NCLX governs superoxide production and OXPHOS function in hypoxia through Na^+^-dependent alteration of inner mitochondrial membrane fluidity. **a-b**, Superoxide detection with DHE in **(a)** siSCR, siRISP, **(b)** non-treated (No treat) or 1 µM myxothiazol (Myxo) treated BAECs. **c, d** Effect of NCLX activation by 10 mM NaCl and 0.1 mM CaCl_2_ on succinate-based OCR in isolated mitochondria from BAECs **(c)**, n=5, or MEFs **(d)**, n=6. **e**, Succinate-cytochrome *c* activity in freeze-thawed mitochondria (i.e. mitochondrial membranes; MM) from BAECs ± 10 mM NaCl (n=4). **f**, Succinate-cytochrome *c* activity in MM from WT or KO MEFs previously subjected to normoxia (Nx) or hypoxia (Hp; 1% O_2_; n=3). **g**, ROS production on succinate-cytochrome c activity in MM from BAECs ± 10 mM NaCl, n=10. **h**, NADH-cytochrome *c* activity in MM from BAECs ± 10 mM NaCl (n=3). **i**, G3P-cytochrome *c* activity in MM from BAECs ± 10 mM NaCl (n=3). **j-k**, Effect of hypoxia (1% O_2_) in Na^+^_mito_ content in WT and KO MEFs, measured with CoroNa-Green adapted for mitochondrial loading and live fluorescence (**j**, n=10), or with SBFI fluorescence after mitochondria isolation (**k**, n=4). **l-m**, FRAP of siSCR or siNCLX BAECs **(l;** n=16) and WT, KO, KO+pNCLX or WT+dnNCLX MEFs **(m;** n=14) expressing mitoRFP in normoxia or hypoxia for 20 min (1% O_2_). **n**, NAO FRAP quench signal of siSCR or siNCLX BAECs and exposed to normoxia or hypoxia for 15 min (1% O_2_; n=5). **o**, Anisotropy of PC liposomes treated with increasing concentrations of NaCl measured by DPH fluorescence (n=3). **p**, Infrared absorption spectra in the region of the carbonyl group of PC liposomes treated or not with 16 mM NaCl. Student’s t-test for pairwise comparisons and one-way ANOVA with Tukey’s test for multiple comparisons. n.s. not significant, * p<0.05, **<p0.01, *** p<0.001. AA = antimycin A. **a,b**, statistical comparisons shown only for Nx vs 0-10 groups.

To date, mitochondrial complexes III (CIII) and I (CI) have been reported to be necessary for ROS production and cellular adaptation to hypoxia^5,6,13^. To examine their contribution, we knocked-down either CI subunit NDUFS4 or CIII subunit RISP, but only NDUFS4 knockdown abrogated Na^+^/Ca^2+^ exchange in hypoxia (Extended Data Fig. 4a-d). Accordingly, pharmacological inhibition of CI, but not of CIII or of complex IV (CIV), blunted hypoxia-induced Na^+^/Ca^2+^ exchange (Extended Data Fig. 4e-f) confirming that CI is necessary for NCLX activation in acute hypoxia.

**Fig. 4.**
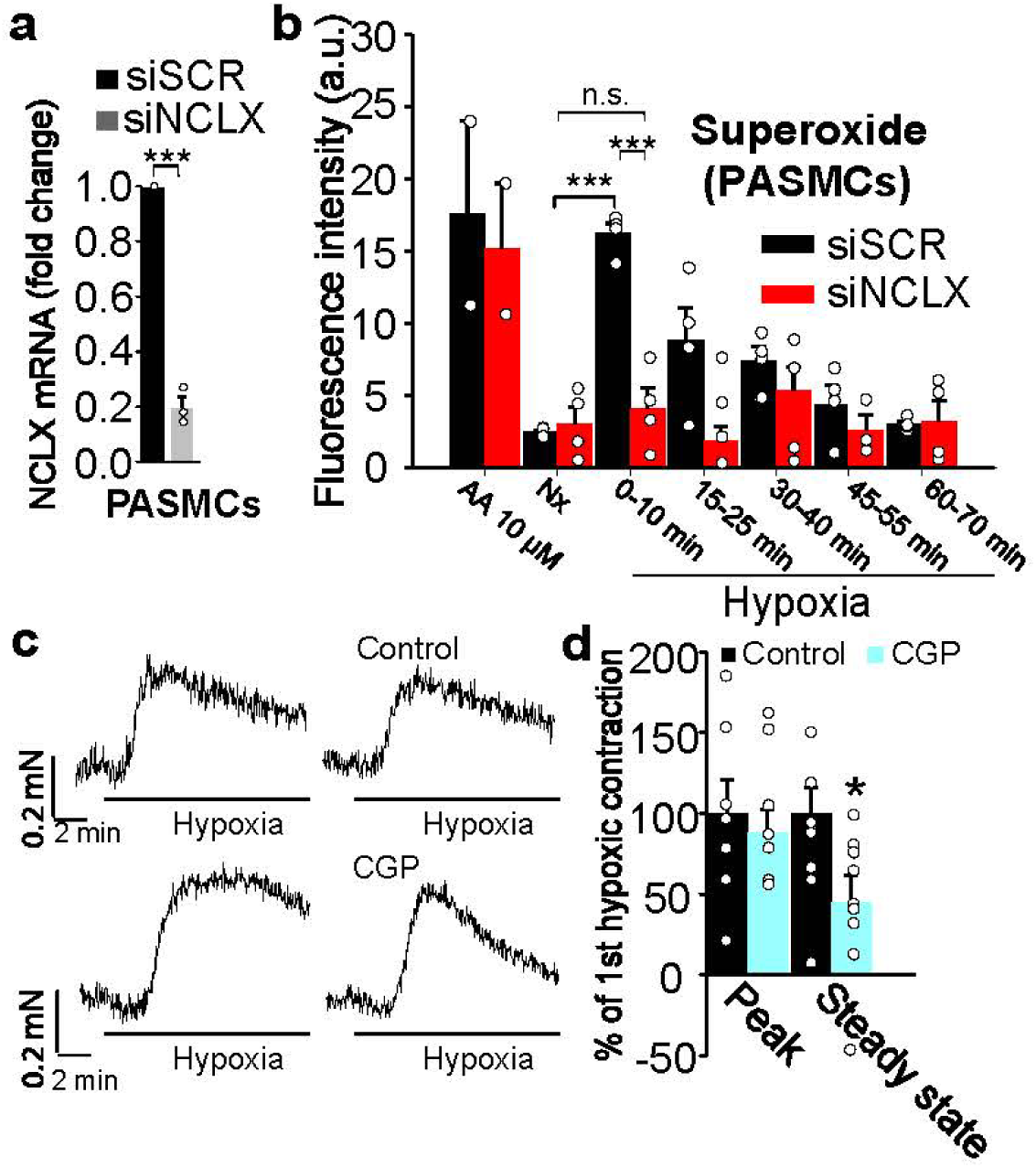
Na^+^_mito_ influx inhibition abolishes hypoxic pulmonary vasoconstriction. **a**, Effectiveness of NCLX silencing in mice pulmonary artery smooth muscle cells (PASMCs) measured by quantitative RT-PCR analysis, n=3. **b**, Superoxide detection by fluorescence microscopy after incubation with DHE in PASMCs treated with siSCR or siNCLX in normoxia (Nx) or hypoxia (Hp; 1% O_2_), n=3. **c-d**, Representative traces **(c)** and average values **(d)** of HPV measured in rat pulmonary arteries. Each artery was exposed twice to hypoxia, and the second hypoxic challenge was performed in the absence or the presence of 30 µM CGP-37157, n=7-9. Student’s t-test for pairwise comparisons and one-way ANOVA with Tukey’s test for multiple comparisons: * p<0.05, **<p0.01, *** p<0.001. **b**, statistical comparisons shown only for Nx vs 0-10 groups.

Next, we evaluated how CI was influencing NCLX activity during hypoxia. CI presents two functionally different conformation states (A-CI and D-CI)^14–16^, and we previously reported that the D-CI conformation, which is related to reduced CI activity^14–16^, is favoured in acute hypoxia^11^. The D-CI conformation exposes Cys-39 of the ND3 subunit^14^ allowing fluorescent labelling at this residue for determination of conformation state^11^. Using this method, we confirmed that five min of hypoxia increased the D-CI conformation and, in parallel, we observed a decrease in the CI reactivation rate, regardless of NCLX inhibition (Extended Data Fig. 5a-d). Previously, we reported that hypoxia promotes a rotenone-sensitive matrix acidification^11^, which we also confirmed is independent of NCLX activity (Extended Data Fig. 5e). In addition, Na^+^/Ca^2+^ exchange was inhibited by rotenone (a CI inhibitor; Extended Data Fig. 5f-g), which further identifies D-CI formation as a necessary step in NCLX activation.

Acute hypoxia increased the matrix Ca^2+^ (Ca^2+^_mito_) concentration and this was augmented by NCLX knock-down (Fig. 2a). Thus, there is an increase in Ca^2+^_mito_ that drives NCLX activation and Ca^2+^ efflux to the cytosol. Intriguingly, BAECs treated with the mitochondrial Ca^2+^ uniporter (MCU) inhibitor Ru360 still displayed an increase in Ca^2+^_cyto_ and Ca^2+^_mito_ (Fig. 2b-c and Extended Data Fig.6a), excluding a role of MCU in this effect. We reasoned that if Ca^2+^ was not being introduced into the mitochondria, its source could be inside the organelles themselves. It has been known for decades that mitochondria harbour electron-dense spots composed by calcium phosphate precipitates which are extremely sensitive to acidification^17^, but whose function remains obscure^18^. We first confirmed by TEM-EDX analysis that the electron-dense spots that we found in the mitochondria were enriched in calcium, regardless of the element used to relativize (Extended Data Fig. 6b). Next, we confirmed that calcium phosphate precipitates decreased in both size and frequency in the presence of FCCP, a drug promoting strong acidification of the mitochondrial matrix, which concomitantly boosted Ca^2+^_mito_ content and elicited NCLX activation (Fig. 2d and Extended Data Fig. 6c-h). Indeed, acute hypoxia also promoted a decrease in the size of the precipitates, which correlated with an increase in soluble Ca^2+^_mito_ concentration (Fig. 2a, d and Extended Data Fig. 6c). Thus, mitochondrial calcium phosphate precipitates are the likely source of NCLX-dependent Ca^2+^_cyto_ increase. We have previously observed that pre-treatment of BAECs with the CI inhibitor rotenone promotes mitochondrial matrix acidification^11^. Since CI is involved in the hypoxic response, it could be related to the solubilization of mitochondrial Ca^2+^ precipitates through matrix acidification. We reasoned that if this was correct, pre-treatment with the CI-inhibitor rotenone should produce by itself matrix acidification and Ca^2+^ increase rendering mitochondria unable to respond to hypoxia. We pre-treated BAECs with rotenone for 30 min and subjected them to hypoxia, maintaining rotenone during the experiment. As hypothesized, 1 µM rotenone blunted the increase in Ca^2+^_mito_ (Fig. 2c). At this point, we concluded that hypoxia induces the D-CI conformation, leading to mitochondrial matrix acidification and solubilization of calcium-phosphate precipitates to liberate Ca^2+^ and activate NCLX.

Next, we tested how NCLX activation triggered mitochondrial ROS production in acute hypoxia. In line with previous reports identifying CIII as a site of ROS production in hypoxia^19,20^, RISP knockdown abolished the superoxide burst in acute hypoxia (Fig. 3a) without affecting NCLX activation (Extended Data Fig. 4b-d). This effect was reproduced by myxothiazol, an inhibitor of CIII Qo site (Fig. 3b). ROS production by active isolated mitochondria transiently increased upon sequential addition of Na^+^ and Ca^2+^, which activates Na^+^ entry and Ca^2+^ extrusion through NCLX^21,22^ (Extended Data Fig. 7a).

The contribution of the different electron transport chain (ETC) complexes to respiration was measured in isolated mitochondria upon addition of different substrates and inhibitors^23^, in the absence or presence of Na^+^ and Ca^2+^ to activate NCLX, with and without NCLX, or in the absence/presence of CGP-37157. In contrast to CIV-dependent respiration, CI- and CII-dependent respirations were decreased upon NaCl and CaCl_2_ addition, but only the reduction of CII-dependent respiration was dependent on NCLX activation (Fig. 3c-d and Extended Data Fig.7b-e). CII+III activity in isolated mitochondrial membranes was unaffected by the addition of Ca^2+^ (Extended Data Fig. 7f), but clearly decreased with Na^+^ addition in a dose-dependent manner (Fig. 3e and Extended Data Fig.7g), which also increased ROS production (Fig. 3g). CII+III activity was decreased in mitochondrial membranes of WT MEFs subjected to hypoxia, an effect abolished by loss of NCLX (Fig. 3f). Dimethylmalonate (DMM, a complex II inhibitor) did not affect NCLX activity, but decreased the superoxide burst during hypoxia (Extended Data Fig. 7h-j). Surprisingly, individual CII or CIII activities showed no variation after Na^+^ addition (Extended Data Fig. 7k-l); neither did CI+III activity (Fig. 3h). G3PD+III activity also decreased in the presence of Na^+^ (Fig. 3i). In summary, while the individual activity of the different ETC complexes is not affected by Na^+^, the combined activities of different electron donors to CoQ with CIII behave differently based on the source of reducing equivalents, FADH_2_ (CII and G3PDH) or NADH (CI).

Electron transfer between CII and CIII (and between G3PD and CIII) is limited by CoQ diffusion through the inner mitochondrial membrane, which depends on its fluidity^24,25^. We assessed whether IMM fluidity was affected by NCLX activation and Na^+^ import during hypoxia. Acute hypoxia markedly increased mitochondrial Na^+^ (Na^+^_mito_) content through NCLX activation (Fig. 3j-k; Extended Data Fig. 7m-p). Employing fluorescence recovery after photobleaching (FRAP) of a red fluorescent protein targeted to the IMM (MitoRFP) we found that NCLX activation in acute hypoxia reduced membrane fluidity; this effect was abolished by NCLX inhibition by siRNA, genetic deletion, overexpression of dnNCLX, or CGP-37157 (Fig. 3l-m, Extended Data Fig 8a and Supplementary videos 1-8). The FRAP signal was restored after ectopic expression of human NCLX in KO cells (Fig. 3m) suggesting direct causation. We also used the lipophilic quenchable fluorescent probe 10-n-nonyl acridine orange (NAO) that binds specifically to cardiolipin^26^, an abundant phospholipid of the IMM^27,28^. The reduced recovery rate of NAO fluorescence after photobleaching confirmed an NCLX-dependent reduction in IMM matrix side fluidity (Fig. 3n and Extended Data Fig. 8b). Reduced CoQ diffusion at the IMM matrix side would slow electron transfer from CIII to oxidized CoQ in the CIII Qi site during Q cycle turnover^29^. Accordingly, we observed that hypoxic NCLX activity promoted an increase in oxidized CoQ_10_ (Extended Data Fig. 8c). The fact that Na^+^ exerts its effect when it is concentrated in the mitochondrial matrix highlights the physiological relevance of the asymmetric distribution of phospholipids between the matrix and the IMS leaflets of the mitochondrial inner membrane IMM^28^.

To further explore the mechanism by which Na^+^ affects IMM fluidity, we measured anisotropy of fluorescent probes in phosphatydilcholine (PC) or PC:cardiolipin liposomes. In this cell free system, we observed that Na^+^ addition (Fig. 3o and Extended Data Fig. 8d-e) reduced the lipid bilayer fluidity. The decrease in membrane fluidity caused by Na^+^ was further supported by ESR experiments^30^, which suggested that Na^+^-PC interaction may occur at the level of phospholipid headgroups (Extended Data Fig. 8f-i). Infrared (IR) spectroscopy of PC liposomes treated with NaCl showed a distinct shift in the absorption peak of the PC’s carbonyl group (Fig. 3p and Extended Data Fig. 8j) and ICP mass spectrometry showed that the stoichiometry of the interaction is 0.29 ± 0.04 (Na^+^:PC). These results demonstrate that Na^+^ specifically interacts with phospholipids such as PC at the level of its carbonyl group, at a ratio of 3PC:1Na^+^, what is in full agreement with previous estimations^31,32^, and suggests that this direct incorporation of Na^+^ is responsible for reduced membrane fluidity during hypoxia.

Altogether, our data demonstrate that acute hypoxia promotes CI A/D transition and mitochondrial matrix acidification^11^, which partially solubilizes mitochondrial calcium phosphate precipitates to increase matrix free Ca^2+^. The increase in matrix Ca^2+^ is then exported to the cytosol in exchange for Na^+^ by NCLX. Matrix Na^+^ directly binds IMM phospholipids and reduces membrane fluidity. This diminishes CoQ diffusion in the IMM matrix side and slows electron transfer between CII and CIII effectively uncoupling the Q cycle of CIII, increasing the half-life of the semiquinone form in the CIII Qo site, and promotes superoxide anion formation^29^.

To study the physiological relevance of this pathway, we investigated if its manipulation altered the response of pulmonary arteries to hypoxia. Rat pulmonary arteries (PA) exposed to hypoxia in the absence of pretone (precontraction) perform a rapid contractile response (peak contraction), which reaches a plateau (steady-state contraction). The latter is dependent on mitochondrial ROS^33–35^. We found that inhibition of mitochondrial Na^+^_mito_ entry by NCLX silencing inhibited the superoxide burst in rat pulmonary artery smooth muscle cells (PASMCs) subjected to acute hypoxia (Fig. 4a-b). Concomitantly, NCLX inhibition markedly reduced the steady-state component of PA hypoxic contraction (Fig. 4 c-d), in a manner similar to that elicited by antioxidants^36^. These observations suggest a physiological role for NCLX in hypoxic pulmonary vasoconstriction (HPV), a phenomenon necessary to match lung ventilation with perfusion, which is crucial for cardiovascular homeostasis.

The role of Na^+^ in the cell has been restricted to the maintenance of membrane potentials and to aid ion exchange at membranes (Na^+^/K^+^, Na^+^/Ca^2+^, etc.). Here we show that Na^+^ directly modulates mitochondrial OXPHOS and ROS signalling with significant implications in tissue homeostasis. In addition, we identify mitochondrial calcium phosphate precipitates as a significant source of mitochondrial soluble Ca^2+^ in physiological conditions.

Interestingly, we noticed that Na^+^ only exerted its effect on CII+CIII or G3PDH+CIII activities, but not on CI+CIII activity. The only model that may explain this selective behaviour is the assembly of the respiratory complexes into supercomplexes. Thus, CoQ_10_ transfer from CI to CIII does not decrease in the presence of Na^+^ because CI+CIII supercomplex activity does not depend on membrane fluidity, as it may utilize the CoQ (CoQ^NADH^) partially trapped in the supercomplex microenvironment^37,38^. This is supported by a recent report describing the structure of the I+III_2_ supercomplex^39^. On the other hand, CoQ transfer from either CII or G3PD to CIII strongly relies on membrane fluidity and consequently, both CII+CIII and G3PD+III (i.e. CoQ^FAD^) activities are lower in the presence of Na^+^. Therefore, the partial segmentation of the CoQ pool into CoQ^NADH^ and CoQ^FAD^ allows the specific CIII-dependent ROS generation upon Na^+^:PC interaction in response to acute hypoxia meanwhile respiration can be maintained by means of supercomplex function.

A recent report described the beneficial role of NCLX by decreasing ROS production in a murine model of cardiac ischemia-reperfusion^12^. Although it could seem contradictory with the findings reported here, both roles of NCLX are compatible. During cardiac ischemia succinate accumulates, which causes an elevation of the CoQH_2_/CoQ ratio and, at reperfusion induces superoxide production by reverse electron transport (RET) through CI^4^. There are three ways by which NCLX activation could hinder ROS production by RET during reperfusion: a) Na^+^:phospholipid interaction would restrain CoQH_2_ diffusion towards CI through the reduction of the IMM fluidity; b) CoQH_2_/CoQ ratio drops during hypoxia in a NCLX-dependent manner, what would decrease RET during reoxygenation; c) Ca^2+^ extrusion by NCLX would prevent further TCA-cycle dehydrogenases activation, limiting the production of reducing equivalents to OXPHOS complexes. Thus, our model fully supports the role of NCLX in attenuating ROS production and injury upon cardiac reperfusion^12^. Indeed, the present mechanism may represent a way by which mitochondria reduce potentially toxic CI RET-derived ROS and induce an adaptive short-time elevation of CIII-dependent ROS production.

In summary, our findings open new horizons for the study of mitochondrial OXPHOS and signalling. The impact of Na^+^ as a second messenger on mitochondrial homeostasis had been ignored, but our results suggest it likely plays a pivotal role in mitochondrial physiology in health and disease.

## Supporting information

Supp Video 1. FRAP MEFs WT Normoxia

Supp Video 2. FRAP MEFs WT Hypoxia

Supp Video 3. FRAP MEFs KO Normoxia

Supp Video 4. FRAP MEFs KO Hypoxia

Supp Video 5. FRAP MEFs KOpNCLX Normoxia

Supp Video 6. FRAP MEFs KOpNCLX Hypoxia

Supp Video 7. FRAP MEFs WTdnNCLX Normoxia

Supp Video 8. FRAP MEFs WTdnNCLX Hypoxia

## Methods

### Animals, cell culture and transfection

All animal experiments were performed following the Guide for the Care and Use of Laboratory Animals and were approved by the institutional ethics committee of the Universidad Autónoma de Madrid or the Universidad Complutense de Madrid, Spain, in accordance with the European Union Directive of 22 September 2010 (2010/63/UE) and with the Spanish Royal Decree of 1 February 2013 (53/2013). All efforts were made to minimize the number of animals used and their suffering.

Cells were routinely maintained in cell culture incubators (95% air, 5% CO_2_ in gas phase, 37 ºC). Bovine aortic endothelial cells (BAECs) were isolated as described^40^ and cultured in RPMI 1640 supplemented with 15% heat-inactivated fetal bovine serum (FBS), 100 U/mL penicillin and 100 μg/mL streptomycin. Mouse embryonic fibroblasts (MEFs) were isolated as described previously^12^ and cultured in DMEM supplemented with 10% heat-inactivated fetal bovine serum (FBS), 100 U/mL penicillin and 100 μg/mL streptomycin. Human umbilical vein endothelial cells (HUVECs) were isolated as described^41^ and cultured in Medium 199 supplemented with 20% heat-inactivated FBS, 16 U/mL heparin, 100 mg/L ECGF (endothelial cell growth factor), 20 mM HEPES (4-(2-hydroxyethyl)piperazine-1-ethanesulfonic acid), 100 U/mL penicillin and 100 μg/mL streptomycin. BAECs were used between passages 3 and 9 and HUVECs between passages 3 and 7. Endothelial morphology was assessed by visual inspection.

Rat pulmonary artery smooth muscle cells (PASMCs) were isolated as described^34^ and cultured in Dulbecco’s modified Eagle’s medium (DMEM) supplemented with 10% FBS, 1.1 g/L pyruvate, 1% non-essential amino acids, 100 µg/mL streptomycin and 100 U/mL penicillin. PASMCs were used between passages 2 and 3.

Hepatoma cell line HepG2 were cultured at 37 ºC in DMEM supplemented with 10% heat-inactivated FBS, 20 mM HEPES, 100 U/mL penicillin and 100 μg/mL streptomycin.

Transfection of 30 nM siRNA or 0.25 µg Cyto-GEM-GECO, Cepia 2mt, human NCLX (pNCLX; Addgene), S468T NCLX (dnNCLX), C199S pHyPer-Myto (mitosypHer), pHyPer-cyto (cytohyper) or pDsRed2-Mito (mitoRFP) vector DNA per 0.8 cm^2^ well was carried out using Lipofectamine 2000 (Invitrogen). Acute ablation in NCLX flx/flx MEFs was obtained by infection with adenoviruses expressing cytomegalovirus (CMV)-Cre (ad-CRE; 300 pfu/cell; Vector Biolabs). In PASMCs, 10 nM siRNA was transfected using Lipofectamine RNA iMAX (Invitrogen); in parallel experiments, siRNA efficiency was measured by RT-PCR using commercially available primers (TaqMan Gene Expression Assay Rn01481405_m1, Applied Biosystems). Experiments were carried out 48 to 72 h after transfection.

Immortalization of HUVECs and MEFs was performed by retrovirus infection. 293T cells were transfected with 12 ug pCL-Ampho and 12 ug pBABE-SV40-puro using Lipofectamine RNA iMAX (Invitrogen). Two days later, 1/2 dilution of 293T cells media and 8 ug/mL of polybrene were added on MEFs or HUVECs cultures during 4 hours. Then, the infection was repeated, but adding 4 ug/mL of polybrene overnight. Next day, the first infection was repeated and, upon 4 hours, MEFs or HUVECs were passaged with 1/1000 of puromicin.

### siRNA preparation

Four double-stranded siRNA against bovine NCLX were designed and purchased from Integrated DNA Technologies and Dharmacon (sense sequences of siNCLX1: AGCGGCCACUCAACUGCCU; siNCLX2: GUUUGGAACUGAAACACU and UUCCGUAAGUGUUUCAGU mixed; siNCLX3: AAAGGUGGAAGUAAUCAC and ACGUAUUGUGAUUACUUC mixed; siNCLX4: GAAUUUGGAGUGAUUCAC and UUUUCAAGUGAAUCACUC mixed). Double-stranded siRNAs against bovine NDUFS4 and RISP were designed and purchased from Integrated DNA Technologies (NDUFS4 sense sequence GCUGCCGUUUCCGUUUCCAAGGUUUTT; NDUFS2 sense sequence TCGGACAGTCGACATTGGGATT; RISP sense sequence CCAAGAAUGUCGUCUCUCAGUUUTT). Scrambled siRNA (siSCR) was purchased from Santa Cruz Biotechnology.

siRNA against rat NCLX (SL8B1 gene) was purchased from OriGene (SLC8B1 Trilencer-27 Rat siRNA).

### Detection of superoxide by fluorescence microscopy in fixed cells

Cells were seeded on glass coverslips one day before experimentation. In some experiments, 1 µM rotenone or 10 µM **7-**chloro-5-(2-chlorophenyl)-1,5-dihydro-4,1-benzothiazepin-2(3*H*)-one (CGP-37157) was added 30 min before experimentation and maintained during the experiment. For treatments in hypoxia, all the solutions were pre-equilibrated in hypoxic conditions before use; plated cells were introduced into an Invivo2 400 workstation (Ruskinn) set at 1% O_2_, 5% CO_2_, 37 ºC, and incubated for the indicated times (0, 15, 30, 45 and 60 min) in fresh medium, washed three times with Hank’s Balanced Salt Solution with Ca^2+^/Mg^2+^ (HBSS+Ca/Mg, which contains 141 mM Na^+^ and 5.8 mM K^+^) and incubated with 5 µM dihydroethidium (DHE) in HBSS+Ca/Mg for 10 min in the dark. Excess probe was removed by three washes with HBSS+Ca/Mg, cells were fixed with 4% paraformaldehyde (PFA), and incubated in the dark at 4 ºC for 15 min. After fixation, the cells were again washed three times with HBSS+Ca/Mg and coverslips were placed on slides. For normoxic treatments, the medium was changed to fresh normoxic medium, and cells were treated as for hypoxic cells but in a standard cell incubator. Images (three images per each coverslip; the number of independent experiments is described in the figure legends) were taken with a Leica DMR fluorescence microscope with a 63× objective, using the 546-12/560 excitation/emission filter pairs, and quantified using ImageJ software (NIH). The same threshold was set for all the images and the mean value from histograms was averaged for the three images of each coverslip.

### Detection of intracellular calcium, sodium, ROS and mitochondrial membrane potential by live imaging fluorescence microscopy

Cells were seeded in 6-well plates one day before experimentation. Plated cells were washed three times with HBSS+Ca/Mg±Glucose and incubated with 30 nM tetramethylrhodamine methyl ester (TMRM), 1 µM Fluo-4 AM, 10 µM CoroNa Green AM or 10 µM 6-carboxy-2’,7’-dichlorodihydrofluorescein diacetate (CDCFDA) for 20 min at 37 ºC in the dark. CDCFDA, CoroNa Green AM and Fluo-4 AM were then washed out and new HBSS+Ca/Mg was added. In some experiments, 10 µM CGP-37157 or 10 µM dimethylmalonate (DMM) were also added and maintained during the remainder of the experiment. For Fluo-4 AM imaging, cells were further incubated for 30 min at 37 ºC in the dark to allow complete de-esterification of the probe. After this time, the plate was placed into a Leica DM 16000B fluorescence microscope equipped with a Leica DFC360FXcamera, an automated stage for live imaging and a thermostated hypoxic cabinet. The planes were focused for image capture, and images were taken with a 20× objective every 2 min during 40 min, providing a total of 21 cycles. Normoxia experiments started and ended at 20% O_2_ and 5% CO_2_, whereas hypoxia experiments started at 20% O_2_ and 5% CO_2_ and then were switched to 2% O_2_ and 5% CO_2_ in cycle 2. The excitation/emission filter pairs used were as follows: and TMRM, and 480-40/505 for CDCFDA, Fluo-4 AM and CoroNa Green AM. Images were quantified with Leica Las-AF software. Three independent experiments were performed for each condition. For each experiment and condition, four regions of interest (ROIs) were created, each ROI surrounding an individual cell, and the mean fluorescence of each ROI for each time cycle was collected. In some analyses, for each experiment and condition, four identical linear ROIs were created and the maximum peak value of cycles 0, 5, 10, 15 and 20 were collected for each ROI.

### Detection of cytosolic calcium, sodium and H_2_O_2_, and intramitochondrial pH, Ca^2+^ and Na^+^ by live imaging confocal microscopy

To detect intracellular calcium and sodium, cells were seeded one day before experimentation, washed three times with HBSS+Ca/Mg+Glucose and in some experiments incubated with 1 µM Fluo-4 AM, Corona Red or 5 µM CoroNa Green AM, Asante Natrium 2 AM (ANG2 AM) for 30 min at 37ºC in the dark. In some experiments, 10 µM CGP-37157, 10 µM DMM, 1 µM FCCP, 1 µM rotenone, or 1 µM Ru360 were also added and maintained during the remainder of the experiment. 100 µM Histamine was added acutely in control experiments. CoroNa Green AM, ANG2 AM, Corona Red and Fluo-4 AM were then washed out and new HBSS+Ca/Mg plus 25 mM glucose was added. For Fluo-4 AM and ANG2 AM imaging, cells were further incubated for 30 min at 37ºC in the dark to allow complete de-esterification of the probe. After this time, the plate was placed into a Leica SP-5 confocal microscope, an automated stage for live imaging and a thermostated hypoxic cabinet. The planes were focused for image capture and images were taken with a 63× objective. In some experiments images were taken every 2 min during 40 min, providing a total of 21 cycles and in others every five minutes. Normoxia experiments started and ended at 20% O_2_ and 5% CO_2_, whereas hypoxia experiments started at 20% O_2_ and 5% CO_2_ and then were switched to 1% O_2_ and 5% CO_2_ in cycle 1. In other experiments hypoxia conditions were set at the middle of the protocol, allowing the quantification of the same cells during normoxia and hypoxia. Loaded cells were excited with an argon/krypton laser using the 496 nm line, except Corona Red which was excited with the 514 line. Fluorescence emission of Fluo-4 AM, ANG2 AM and CoroNa Green was detected in 515–575 nm range, and 555-575 nm for Corona Red.

To detect intramitochondrial pH, cells were transfected with the ratiometric probe mitosypHer in 8-well plates the day before the experiment. The same protocol as above was used, except that the objective was 63× and imaging time was 30 min, with 7 cycles of 5 min. Excitation was performed with a 405 diode laser for 405 nm line and an argon/krypton laser for 488 nm line, and fluorescent emission was recorded at the 515–535 nm range.

For H_2_O_2_ detection, cells were transfected with the non-targeted version of HyPer (pHyPer-cyto) following the same procedure for live imaging as with mitosypHer.

Images were quantified with ImageJ software. Three or four independent experiments were performed for each condition. For each experiment and condition in loaded cells, four identical linear regions of interest (ROIs) were quantified, and for each time point the mean of these ROIs was obtained.

For cytosolic Ca^2+^ detection, cells were transfected with the non-targeted version of GEM-GECO (cyto-GEM-GECO) in 8-well plates the day before the experiment. The objective was 63× and imaging time was 45 min, with 9 cycles of 5 min. Excitation was performed with a 405-diode laser for 405 nm line, and fluorescent emission was recorded at 460 and 510 nm lines.

For mitochondrial Ca^2+^ detection, cells were transfected with the mitochondrial-targeted Cepia2mt or mito-GEM-GECO in 8-well plates the day before the experiment. The objective was 63× and imaging time was 40 min, with 20 cycles of 2 min. Samples transfected with Cepia2mt were excited with an argon/krypton laser using the 488 nm line and fluorescence emission was detected in the 515–535 nm range.

For mitochondrial Na^+^ detection, cells were washed three times with HBSS+Ca/Mg+Glucose, incubated for 1 hour with 10 µM CoroNa Green AM, washed three times again with HBSS+Ca/Mg+Glucose and incubated a further hour in HBSS+Ca/Mg+Glucose. During this period Corona Green is actively pumped out from the cytosol. Then, after washing three more times with HBSS+Ca/Mg+Glucose, cells were placed in the microscope. The objective was 63× and imaging time was 25 min, with 15 cycles of 2.5 min. Samples were excited with an argon/krypton laser using the 488 nm line and fluorescence emission was detected in the 500–575 nm range.

Calibrations in Ca^2+^ measurements were performed as previously described^42^.

### Western blot analysis

Protein samples were extracted with non-reducing Laemmli buffer without bromophenol blue and quantified by Bradford assay. Extracts were loaded onto 10% standard polyacrylamide gel electrophoresis after adding 5% 2-mercaptoethanol, and subsequently transferred to nitrocellulose membranes or PVDF membranes. The following antibodies were used: polyclonal anti-NCLX antibodies (ab136975, Abcam; ARP44042_P050, Aviva Systems Biology), monoclonal anti-Fp70 (459200; Invitrogen), monoclonal anti-NDUFS4 antibody (ab87399; Abcam), monoclonal anti-RISP (UQCRFS1) antibody (ab14746; Abcam), and monoclonal anti-α-tubulin antibody (#T6199, Sigma). Antibody binding was detected by chemiluminescence with species-specific secondary antibodies labelled with horseradish peroxidase (HRP), and visualized on a digital luminescent image analyser (Fujifilm LAS-4000), with the exception of ab136975 and 459200 which were detected by fluorescence as previously described^43^.

### Quantitative Real Time PCR

Total RNA was extracted from PASMCs using Trizol reagent (Vitro) and 0.5 µg was reverse-transcribed (Gene Amp Gold RNA PCR Core Kit; Applied Biosystems). PCR was performed with GotaqqPCR Master Mix (Promega) with 1 µL of cDNA and specific primer pairs (Rn01481405_m1). β-actin mRNA was measured as an internal sample control.

### Measurement of cellular oxygen consumption

Oxygen consumption rate (OCR) was measured using an XF24 Extracellular Flux Analyzer (Seahorse Bioscience, North Billerica, MA, USA). BAECs, 6×10^4^ per well (6–7 wells per treatment for each independent experiment) were plated one day before the experiment. Cells were preincubated with unbuffered DMEM supplemented with 25 mM glucose, 1 mM pyruvate, and 2 mM glutamine for 1 h at 37 °C in an incubator without CO_2_ regulation. OCR measurements were programmed with successive injections of unbuffered DMEM, 5 µg/mL oligomycin, 300 nM carbonyl cyanide 4-(trifluoromethoxy) phenylhydrazone (FCCP), and 1 µM rotenone plus 1 µM antimycin A. DMSO or 10 µM CGP-37157 were added before starting the measurements. In experiments using silenced cells, as proliferation and cell growth may vary after plating, protein concentration was quantified by the BCA assay to normalize the OCR. Calculations were performed following the manufacturer’s instructions. After measuring basal respiration, oligomycin was added to inhibit respiration (by blocking H^+^-ATPase); therefore, the amount of oxygen used to produce ATP by OXPHOS is estimated from the difference with basal oxygen consumption (i.e., coupling efficiency). FCCP uncouples OXPHOS by translocating H^+^ from intermembrane space to matrix, thus maximizing electron flux through the electron transport chain (ETC), giving the maximal respiration rate. This treatment provides information about the stored energy in mitochondria that a cell could use in an energetic crisis (i.e., reserve capacity). Antimycin A and rotenone block complexes III and I, respectively, consequently inhibiting electron flux through the ETC and eliminating any H^+^ translocation, therefore, the leftover value is non-mitochondrial respiration, that is, the oxygen consumed by other enzymes in the cell.

### Fluorescent labelling of ND3 Cys-39 from isolated mitochondrial membranes

SMP were prepared as previously described ^11^. SMP protein amount was determined by BCA assay and then proteins were solubilized with 4 g/g digitonin, incubated 5 min on ice and centrifuged for 30 min at 16,000 g, 4 ºC. Samples were split into two parts: one part was incubated at 37 ºC for 60 min to fully deactivate complex I and the other part was kept on ice. Samples were then incubated with Bodipy-TMR C5-maleimide (Invitrogen) for 20 min at 15 ºC in the dark; then, 1 mM cysteine was added and the samples were further incubated for 5 min. After this time, the samples were precipitated twice with acetone, centrifuged at 9,500 g for 10 min at 4 ºC in the dark, and the resulting pellet was resuspended in non-reducing Laemmli loading buffer. For each sample, 100 µg were loaded onto 10% Tricine-SDS-PAGE gels as described^44^. Total protein staining was performed with Sypro Ruby (Invitrogen). The images of the different fluorophores were obtained using a digital fluorescent image analyzer (Fujifilm LAS-4000). Images were quantified using ImageQuant TL7.0 software.

### Mitochondria isolation and measurement of H_2_O_2_ and oxygen consumption

Mitochondria were isolated from BAECs with a protocol adapted for cell culture^37^. Briefly, after resuspending BAECs with a sucrose buffer in a glass Elvehjem potter, homogenization was performed by up and down strokes using a motor-driven Teflon pestle. Successive homogenization-centrifugation steps yielded the mitochondria-containing fraction.

H_2_O_2_ production from isolated rat heart mitochondria was performed in an O2k Oxygraph instrument (Oroboros Instruments). 500 µg of isolated rat heart mitochondria were loaded in KCl buffer with Amplex Red: 150 mM KCl, 10 mM K_2_HPO_4_, 1 mM EDTA, 5 mM MgCl_2_, 1 mg/mL BSA,15 µg/ml Horseradish peroxidase (HRP), 15 µg/ml superoxide dismutase (SOD) and 25 µM Amplex Red. Recordings started before the addition of substrates: 5 mM glutamate and 5 mM malate (GM); and the indicated amounts of NaCl and/or CaCl_2_ were subsequently added.

Oxygen consumption was determined with an oxytherm Clark-type electrode (Hansatech) as described^37^. Briefly, isolated mitochondria from BAECs (150 µg), MEFs WT or MEFs KO were resuspended in MAITE buffer (10 mM Tris-HCl, pH 7.4; 25 mM sucrose; 75 mM sorbitol; 100 mM KCl; 10 mM K_2_HPO_4_; 0.05 mM EDTA; 5 mM MgCl_2_; 1 mg/mL BSA) either untreated, treated with 20 mM NaCl and 100 µM CaCl_2_ or with 20 mM NaCl, 100 µM CaCl_2_ and 10 µM CGP-37157. Substrates and inhibitors were then successively added: 5 mM glutamate and 5 mM malate (GM), 1 µM rotenone, 10 mM succinate, 2.5 µg/mL antimycin A, 10 mM N,N,N’,N’-tetramethyl-p-phenylenediamine and 10 mM sodium azide. Oxygen consumption rate (OCR) was obtained by calculating the slope after each treatment. Values for specific complex input were acquired from the subtraction of substrate less specific inhibitor rates.

### Mitochondrial membranes isolation and complex activity measurement

Mitochondrial membranes from BAECs were obtained after freezing-thawing isolated mitochondria, and OXPHOS enzyme activity was measured as described^37^, using 50 µg per sample. Briefly, rotenone-sensitive NADH-ubiquinone Q_1_ oxidoreduction (complex I activity) was measured by changes in absorbance at 340 nm. Succinate dehydrogenase (complex II) activity was recorded in a buffer containing mitochondrial membranes, succinate and 2,6-dichlorophenol-indophenol (DCPIP) by changes in absorbance at 600 nm. Rotenone-sensitive NADH-cytochrome *c* activity (complex I+III activity) was measured by changes in absorbance at 550 nm after NADH addition. Antimycin A-sensitive succinate-cytochrome *c* activity (complex II+III activity) was calculated after measuring changes in absorbance at 550 nm. ROS production in CII+CIII activity was monitored by fluorescence of dichlorofluorescein probe (DCF). Antimycin A-sensitive ubiquinone 2-cytochrome *c* activity (complex III activity) was measured following changes in absorbance at 550 nm. Glycerol-3-Phosphate dehydrogenase (G3PD)+III activity was measured as for CII+III activity but with glycerol-3-Phosphate (10 mM) as electron donor.

### Measurement of mitochondrial Na^+^

In some experiments, cells were preincubated with 10 µM CGP-37157 for 30 min. Then, cells were treated with normoxia, or hypoxia (1%O_2_) for 10 min. Mitochondria were isolated on ice and resuspended in milliQ water. The samples were split, one part incubated with benzofuran isophthalate tetra-ammonium salt (SBFI) and the other was used to quantify protein amount by BCA. For SBFI measurements, a calibration curve was used in every measurement and the fluorescence recorded at 340 nm/380 nm and emission 520 nm in a FLUOstar Omega Microplate Reader (BMG Labtech).

### Fluorescence recovery after photobleaching (FRAP)

Cells were transfected with pDsRed2-Mito Vector (Clontech). Growing media was changed for HBSS+Ca/Mg+Glucose ± 10 µM CGP-37157 and the plate placed into a Leica SP-5 confocal microscope, an automated stage for live imaging and a thermostated hypoxic cabinet. The planes were focused for image capture and images were taken with a 63× objective with 13x zoom.

Samples were excited with an argon/krypton laser using the 514 nm line and emission was detected in the 565–595 nm range. Images were collected using TCS software (Leica). MitoRFP was scanned five times and then bleached using 15 scans at 40% laser power. To image the recovery of fluorescence intensity after photobleaching, we recorded 60 scans every 1 s. FRAP in normoxia was performed at 20% O_2_ and 5% CO_2_, the chamber was then switched to 1% O_2_ and 5% CO_2_ and after 25 min FRAP was performed in hypoxia.

### Fluorescence quenching

Medium of BAECs was changed for HBSS+Ca/Mg+Glucose with 50 nM 10N-nonyl acridine orange (NAO). In some experiments 10 µM CGP-37157 was added. The plate placed into a Leica SP-5 confocal microscope, an automated stage for live imaging and a thermostated hypoxic cabinet. The planes were focused for image capture and images were taken with a 63× objective with 13x zoom.

Samples were excited with an argon/krypton laser using the 488 nm line and emission was detected in the 515–535 nm range. Images were collected using TCS software (Leica). NAO was scanned two times and then bleached using 15 scans at 20% laser power. To image the quench of fluorescence intensity after photobleaching, we recorded 30 scans every 0.372 s. Quenching recording in normoxia was performed at 20% O2 and 5% CO2, the chamber was then switched to 1% O2 and 5% CO2 and the quench again recorded after 15 min in hypoxia.

### Fourier-transform infrared spectroscopy (FT-IR)

Phosphatidylcholine liposomes were prepared using the thin film hydration method followed by extrusion with filters of decreasing diameter (400, 200 and 100 nm)^45^. Lipids were hydrated with water with minimal metal impurities (Optima™ LC/MS Grade, Fisher Chemical). Liposomes were concentrated by filtration up to 72 mg/mL of lipid content. Lipid concentration was determined through the Rouser assay^46^. Sodium salt was mixed and incubated at 37 ºC (2h) with the liposomes at a ratio of 16 mM X+:0.5 mg/mL lipids. FT-IR spectra of liposomes and liposomes incubated with alkali metals were collected afterwards with a Nicolet 6700 FT-IR spectrometer (Thermo Scientific) in transmission mode in liquid samples. All experiments were performed in triplicate.

### Inductively-Coupled Plasma Mass Spectroscopy (ICP-MS)

Phosphatidylcholine liposomes were mixed and incubated at 37 ºC (2h) with sodium chloride salt at a ratio of 16 mM X+:0.5 mg/mL lipids. Thereafter, solutions were washed by filtration with Optima™ LC/MS Grade water and digested with HNO3 acid ([Na+]= 13 ppt, Nitric Acid Optima, for Ultra Trace Elemental Analysis, Fisher Chemicals). The number of alkali cations bound to each lipid within the liposomes, after 3 washing steps, was determined with ICP-MS iCAP-Q from Thermo Scientific equipped with a collision/reaction cell and Kinetic Energy Discrimination (KED). Lipid concentration was determined through the Rouser assay^46^. All experiments were performed in triplicate.

### Transmission electron microscopy

EM of cells and isolated mitochondria were performed as described^47^ and thin sections were imaged on a Tecnai-20 electron microscope (Philips-FEI). Quantification of area and frequency of electron dense spots was performed manually and using Image J software.

### Transmission electron microscopy and energy-dispersive X-ray spectroscopy analysis (TEM-EDX)

Cell samples were fixed using 4% formaldehyde (EMS) and 2.5% glutaraldehyde (Sigma) in 0.1 M HEPES for 4 hours at 4ºC. Samples were post-fixed 1 hour at RT in a 2% osmium tetroxide (EMS) and 3% potassium ferrocyanide (Sigma) solution. Cell preparations were dehydrated through graded acetone series and embedded in Spurr’s Low Viscosity embedding mixture (EMS). Ultrathin sections (70 nm) were then mounted on formvar coated copper grids and stained with lead citrate. Samples were examined on a Jeol JEM 1400 with SDD (Silicon Drift Detector) and elemental content estimation was performed under 80.000-100.000 magnification and using 50-100 seconds lifetime acquisition. Data on oxygen (O), carbon (C), lead (Pb) and calcium (Ca) were collected from the TEM-EDX analysis. C, O and Pb were used as controls.

### Measurement of redox state of ubiquinone

The ratio of the oxidized and reduced forms of ubiquinone (CoQ) were measured as described^48–50^. 10 µM CGP-37157 or antimycin A was added 30 min before the experiment and maintained throughout. HUVECs with or without CGP-37157 were subjected to 10 min of hypoxia (1% O_2_) or normoxia. Antmicyin A was used in normoxic cells as a control. Plates were washed three times with ice-cold PBS and the cells were transferred to ice-cold tubes, which were subsequently centrifuged at 2000 g for 5 min at 4º C. Cell pellets were resuspended in 95 µl of PBS from which 91.5 µl were taken and mixed with 3.5 µl 14.3 µM 2-mercaptoethanol. 5 µl 10 µM coenzyme Q6 were added as internal standard. Then, 330 µl 1-propanol were added, the sample vortexed for 30 s, incubated for 3 min at RT, vortexed again for 20 s and centrifuged at 14,000 g for 5 min. 100 μl of the supernatant was immediately injected into a 166-126 HPLC system (Beckman-Coulter) equipped with an UV/Vis detector (System Gold R 168, Beckman-Coulter) and an electrochemical (Coulochem III ESA) detector. Separation was carried out in a 15 cm Kromasil C18 column (Scharlab, Spain) at 40ºC with a mobile phase 20 mM ammonium acetate pH 4.4 in methanol (solvent A) and 20 mM ammonium acetate pH 4.4 in propanol (solvent B). A gradient method was used with a 85:15 solvent mixture (A:B ratio), flow rate of 1.2 mL/min, as the starting conditions. The mobile phase turns to a 50:50 A:B ratio starting at minute 6 and completed in minute 8, flow rate decreases in parallel to 1.1 mL/min. After 20 min (run time), the columns are re-equilibrated to the initial conditions for three additional minutes. UV-spectrum was used to identify the different forms of ubiquinone (oxidized CoQ10 with maximum absorption at 275 nm) and ubiquinol (reduced CoQ, CoQ_10_H_2_, with maximum absorption at 290 nm) using specific standards. Quantification was carried out with electrochemical detector reading s(channel 1 set to −700 mV and channel 2 set to +500 mV, conditioning guard cell after injection valve).

### Electron spin resonance (ESR) of lipid vesicles

ESR spectroscopy is particularly useful to investigate the rotational dynamics of labeled molecules in solutions or membranes. The rotational dynamics timescale, τ_C_, is inversely related to the fluidity of the microenvirement of the probe. ESR experiments were carried out with a 9.5GHz ESR Bruker spectrometer (RSE Bruker EMX). 2 mM DOPC unilamelar vesicles were obtained by ultrasonication. The spin labels (0.25% mol of total lipid composition) were added to the dry lipids before vesicle formation to obtain a symmetrical distribution of probes in the two leaflets. We used 5-doxyl palmitoyl PC (5-Doxyl PC), 12-doxyl palmitoyl PC (12-Doxyl PC) and 16-doxyl palmitoyl PC (16-Doxyl PC). The different position of the probe along the alkyl chain determines the local motional profiles in the three main regions of the lipidic bilayer, near the polar head group (5-Doxyl PC), at the middle region of the hydrophobic chain (12-Doxyl PC) or at the end of the hydrophobic chain (16-Doxyl PC). The ESR spectrum of 5-Doxyl PC incorporated into the lipidic membranes made of DOPC shows an anisotropic (slow) motion whereas those of both 12-Doxyl PC and 16-Doxyl PC reflects an isotropic motion of the acyl-chain. The τ_C_ of 5-Doxyl PC is proportional to the outermost separation between the spectral extrema, 2A_max_, whereas the τc of 12-Doxyl PC and 16-Doxyl PC were calculated by the semiempirical formula established^51^, 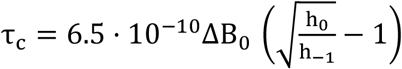 where ∆B_0_ is the width of the central line in Gauss (G), and h_0_ and h_-1_ are the heights of the mid- and high-field lines respectively. 2A_max_, ∆B_0_, h_0_and h_-1_were obtained from the first derivative of each absorption spectrum.

### Steady-state fluorescence emission anisotropy of DPH or TMA-DPH incorporated into lipid vesicles

Mixtures of 1-3 mg of egg phosphatidylcholine (PC; Avanti Polar Lipids Inc., Birmingham, AL, USA), cardiolipin (CL; Avanti Polar Lipids Inc., Birmingham, AL, USA) to a final molar PC:CL molar ratio of 4:1 (when present), and the fluorescent lipid probes diphenyl-1,3,5-hexatriene (DPH; Sigma-Aldrich) or 1-(4(trimetylamonium)-phenyl)-6-diphenyl-1,3,5-hexatriene (TMA-DPH; Sigma-Aldrich), at a lipid/probe molar ratio of 1/100, were dissolved in 200 µL of chloroform:methanol (2:1) and dried under flow of nitrogen for at least 60 min. The resulting dry lipid film was then resuspended in 1.5 mL of MilliQ-degree water and vortexed vigorously for 1 min. This suspension of multilamelar vesicles was then incubated for one hour at 37 ºC. Large unilamelar vesicles were then prepared by extrusion of this lipid dispersion through polycarbonate filters of controlled 100 nm diameter pore size. These vesicles were maintained at 37 ºC for no more than 3 h before being used ^52^. The steady-state fluorescence emission anisotropy experiments were carried out on a SLM Aminco 8000C spectrofluorimeter (SLM Aminco, Urbana, IL, USA), as previously described ^53^. Fluorescence anisotropy (r) of DPH or TMA-DPH incorporated into the liposomes described above was recorded at 37 ºC using excitation and emission wavelengths of 350 or 356 and 452 or 451 nm, respectively.

### Measurement of pulmonary artery contraction

Third division branches of the pulmonary arteries (PAs) were isolated from male Wistar rats and mounted in a wire myograph. Contractile responses were recorded as reported ^50^. The chambers were filled with Krebs buffer containing (in mM) NaCl 118, KCl 4.75, NaHCO_3_ 25, MgSO_4_ 1.2, CaCl_2_ 2.0, KH_2_PO_4_ 1.2 and glucose 11, maintained at 37 °C and aerated with 21% O_2_, 5% CO_2_, 74% N_2_ gas (pO_2_ 17–19 kPa). After an equilibration period of 30 min, PAs (internal diameter 300–400 µm) were distended to a resting tension corresponding to a transmural pressure of 2.66 kPa. Preparations were initially stimulated by raising the K^+^ concentration of the buffer (to 80 mM) in exchange for Na^+^. Vessels were washed three times and allowed to recover. Then, each vessel was exposed to two hypoxic challenges (95% N_2_, 5% CO_2_; pO_2_ = 2.6–3.3 kPa), the second one after 40 min incubation with vehicle (control) or CGP-37157 (30 µM).

### Statistics

Data are presented as mean ± SEM. Normality and homoscedasticity tests were carried out before applying parametric tests. For comparison of multiple groups, we performed one-way analysis of variance (ANOVA) followed by Tukey’s or Bonferroni’s tests for all the groups of the experiment. For comparison of two groups, we used Student’s two-tailed t-test; when the data did not pass the normality test, we used a non-parametric t-test (Mann-Whitney U test). Linear correlation was estimated calculating the Pearson correlation coefficient. Variations were considered significant when the p value was less than 0.05. Statistical analysis was performed with SigmaPlot 11.0 or GraphPad Prism 8 software.

### Data availability

The data that support the findings of this study are available from the corresponding author upon reasonable request.

## Acknowledgments

We thank Dr. Mariusz Kowalewski (Institute of Veterinary Anatomy, UZH) for allowing us the use of the microscope for live cell imaging. We thank Alejandra Alfuzzi, Javier Prieto, Alejandro Mellado (IIS-IP) for collaboration in experiments, Esther Fuertes-Yebra (IIS-IP), for technical assistance, Dr. María Eugenia Soriano and Dr. Federico Caicci (Univ. Padova) for performing electron microscopy, Dr. Judith Langer from CIC biomaGUNE for fruitful discussion and support with the IR spectroscopy measurements, Dr. María Cano and Prof. Antonio G. García (IIS-IP and UAM), Dr. Carlos Rueda and Prof. Jorgina Satrústegui (CMBSO, UAM-CSIC), Prof. Mike Murphy (MRC & Univ. Cambridge), Dr. Israel Sekler (Ben-Gurion University), Dr. Ilka Wittig (Goethe Universität), Dr. José Miguel Mancheño (IQFR, CSIC) and Dr. Alberto Pascual and Prof. José López-Barneo (IBIS, US-CSIC) for helpful discussions, and Dr. Luis del Peso (UAM) and Prof. Francisco Sánchez-Madrid (IIS-IP and UAM) for their support. This research has been financed by Spanish Government grants (partially funded by the European Union FEDER/ERDF) CSD2007-00020 (RosasNet, Consolider-Ingenio 2010 programme to A.M.R. and J.A.E.), CP07/00143, PS09/00101, PI12/00875, PI15/00107, and RTI2018-094203-B-I00 (to A.M.R), CP12/03304 and PI15/01100 (to L.M.), SAF2016-77222-R (to A.C.), SAF2015-65633-R (to J.A.E), SAF2013-32223 (to M.G.L.) and SAF2017-84494-2-R (to J.R.C.), by the European Union (ITN GA317433 to J.A.E. and MC-CIG GA304217 to R.A.P.), by a grant from the Comunidad de Madrid (B2017/BMD-3727, to A.C.), by a grant from the Fundación Domingo Martínez (to M.G.L. and A.M.R.), by grants from the Fundación BBVA (to R.A.P. and J.R.C.), by the Programa Red Guipuzcoana de Ciencia, Tecnología e Información 2018-CIEN-000058-01 (J.R.C) and from the Basque Government under the ELKARTEK Program (Grant No. KK-2019/bmG19 to J.RC.), by Swiss National Science Foundation (SNF) grant 310030_124970/1 to A.B, by a travel grant from the Instituto de Investigación Sanitaria Princesa (to P.H.-A.) and by the COST actions TD0901 (HypoxiaNet) and BM1203 (EU-ROS). CNIC is supported by the Pro-CNIC Foundation and is a Severo Ochoa Center of Excellence (Spanish Government award SEV-2015-0505). CIC biomaGUNE is supported by the María de Maeztu Units of Excellence Program from the Spanish Government (MDM-2017-0720). P.H.-A. was recipient of a predoctoral FPU fellowship from the Spanish Government, E.N. is recipient of a predoctoral FPI fellowship from the Universidad Autónoma de Madrid (UAM), and A.M.-R., L.M. and J.E. are supported by the I3SNS or “Miguel Servet” programmes (ISCIII, Spanish Government; partially funded by FEDER/ERDF).

## Author contributions

P.H.A. and A.M.R. conceived the idea. P.H.A., J.A.E. and A.M.R. designed the study. P.H.A, C.C.F. E.R. and T.O. performed the bulk of the experiments and analysed the data. A.V.L.V. performed EDX experiments. L.M. and A.C. carried out PA contractility experiments. P.H.A., A.C.R., J.C.R.A. and P.N. analysed the redox state of ubiquinone. P.H.A., R.A.P. and J.A.E. analysed mitochondrial complex and supercomplex activities. E.N., E.P., A.P.A., J.E., M.G.L., E.R., J.D.C.G., T.V.P., A.I.A., D.T., P.J. and J.W.E. helped with crucial experimental procedures. P.J. and J.W.E. provided NCLX KO fibroblasts. I.L.R. performed ESR experiments with liposomes. A.M.P. performed experiments with DPH or TMA-DPH labelled liposomes, P.H.A., S.C.R. and J.R.C. performed IR and ICP-MS experiments of Na^+^-phospholipid interaction. A.B., J.A.E. and A.M.R. supervised the study and P.H.A., J.A.E. and A.M.R. wrote the manuscript.

The authors declare no competing financial interest.

## Extended Data

**Extended Data Fig. 1.**
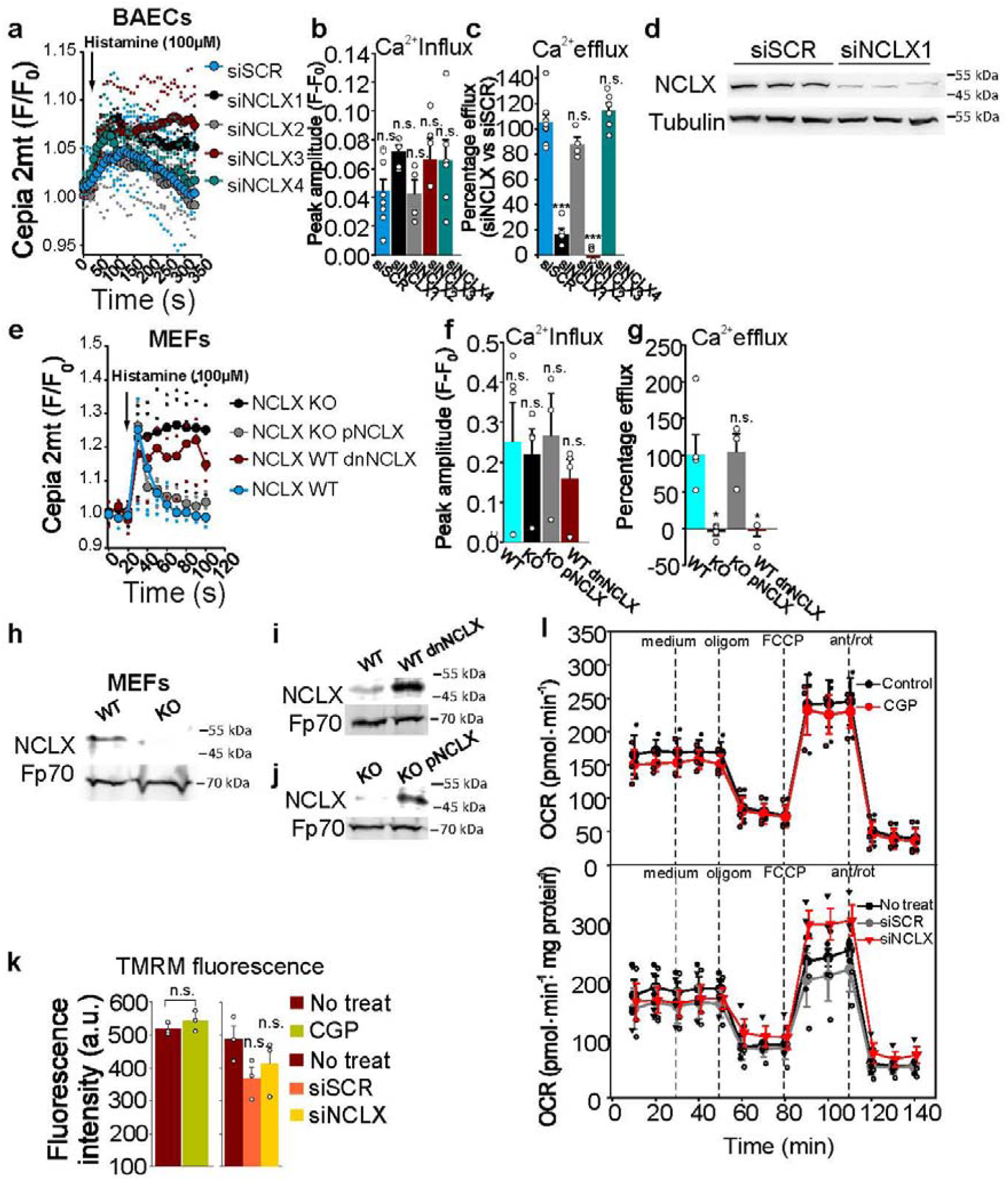
Interference of *Slc8b1* affects NCLX function, but inhibition of NCLX has no effect on general mitochondrial function. **a-c**, Assessment of the effect of different siRNAs against *Slc8b1* (siNCLX) on mitochondrial Ca^2+^ influx and efflux rates in BAECs transfected with the mitochondria-directed Ca^2+^ reporter protein Cepia 2mt (n≥4), after addition of histamine; **a**, mean traces plotted as fluorescence change relative to initial fluorescence (F/F_0_); **b**, Mitochondrial Ca^2+^ peak amplitude (calculated from the highest value of fluorescence minus the first basal value of fluorescence).; **c**, Percentage of mitochondrial Ca^2+^ efflux (calculated from the highest value of fluorescence minus the lowest value of fluorescence after histamine application, relative to percental siSCR values). **d**, Assessment of NCLX protein amount after interference in whole-cell extracts from BAECs, by Western blot (representative image of n=3).. **e-g**, Mitochondrial Ca^2+^ influx and efflux rates in WT, KO, KO+pNCLX or WT+dnNCLX MEFs transfected with Cepia 2mt (n≥4) after addition of histamine, as in **a-c**. **h-j**, Assessment of NCLX protein amount in MEFs mitochondrial extracts by Western blot (representative images of n=3). **k**, Mitochondrial membrane potential in BAECs measured with 20 nM TMRM in non-quenching mode (n=3). **l**, Oxygen consumption rate (OCR) in BAECs with NCLX inhibition by 10 µM CGP-37157 (upper panel, n=3), or NCLX interference with siNCLX1 (lower panel, n=4).

**Extended Data Fig. 2.**
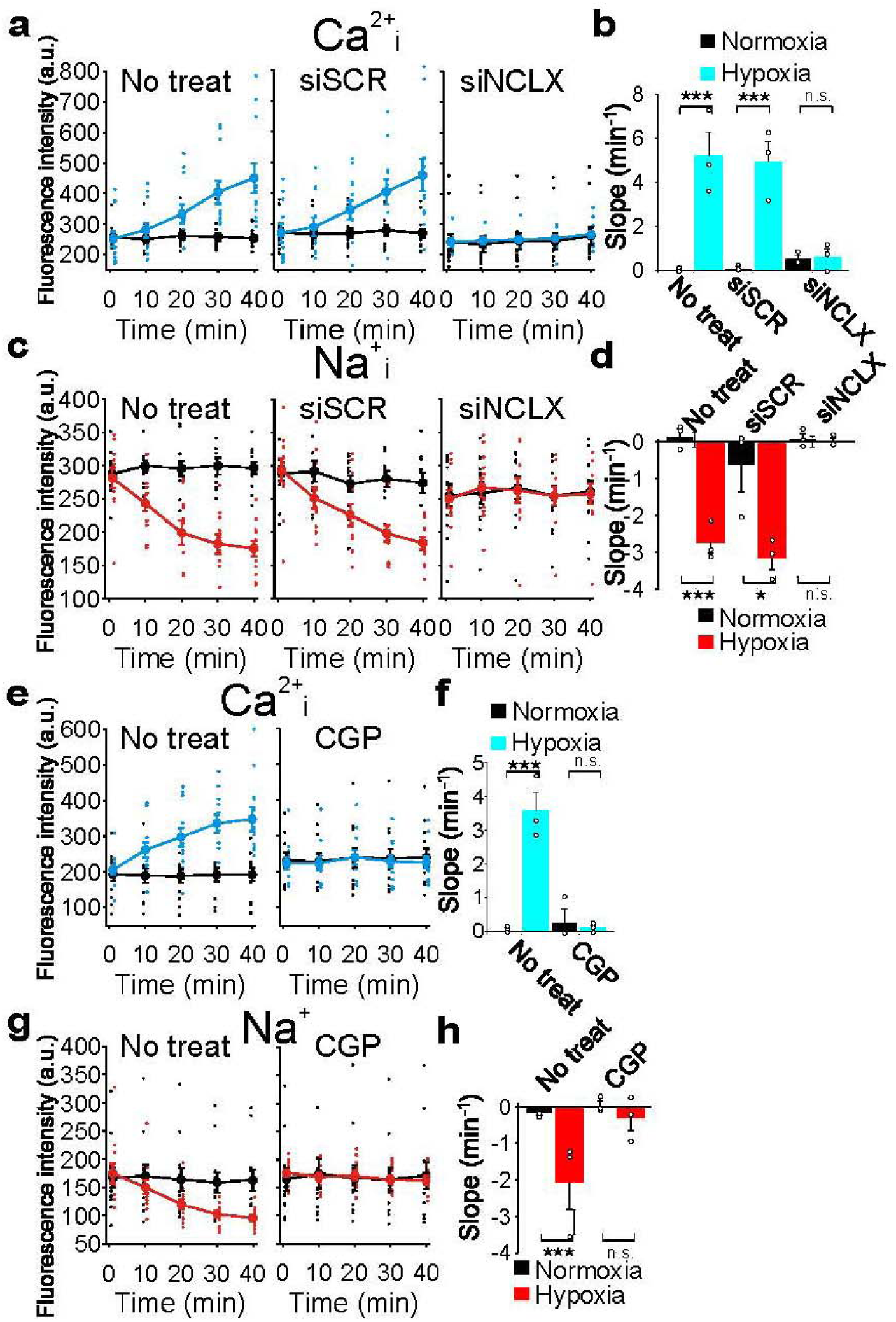
Hypoxia activates Na^+^/Ca^2+^ exchange through NCLX. Cytosolic Ca^2+^ (Ca^2+^_i_) or Na^+^ (Na^+^_i_) were measured by live imaging fluorescence microscopy with Fluo-4 AM or CoroNa Green, respectively, in normoxia or acute ischemia (2% O_2_ without glucose). **a-d**, BAECs not treated (No treat) or transfected with siSCR or siNCLX. **e-h**, BAECs treated or not with the NCLX inhibitor CGP-37157 (10 µM). **a,c,e,g**, time-course traces; **b,d,f,h**, slopes. n=3. * p < 0.05, ** p < 0.01, *** p<0.001, n.s. not significant (Student’s t-test).

**Extended Data Fig. 3.**
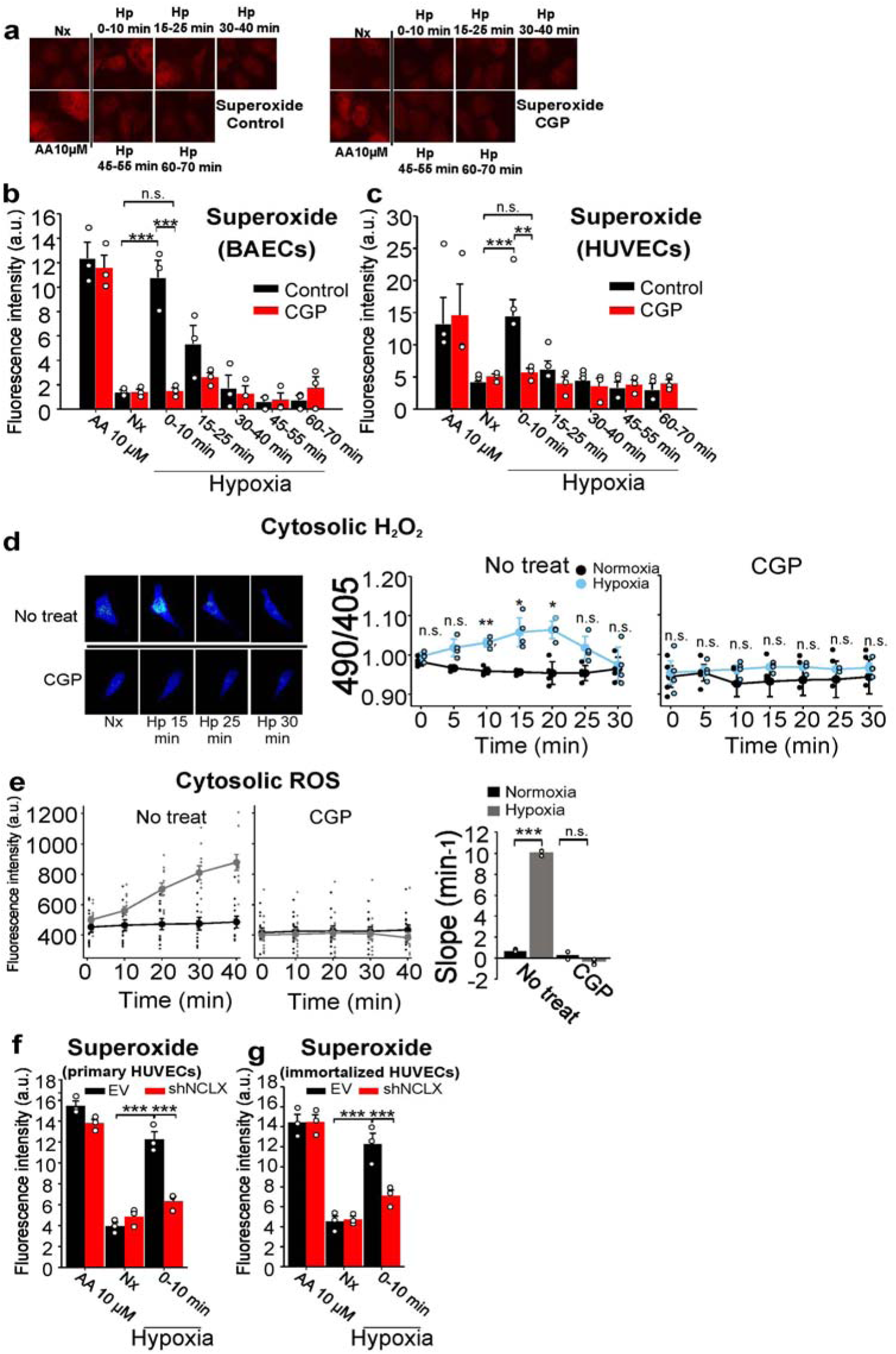
Inhibition of NCLX prevents the increase in ROS production triggered by hypoxia. **a–c**, Superoxide detection by fluorescence microscopy after incubation with DHE in 10-min time windows in normoxia (Nx) or hypoxia (1% O_2_); AA = antimycin A. **a-b**, BAECs with 10 µM CGP-37157, images of one experiment and mean intensity of three independent experiments; **c**, HUVECs with 10 µM CGP-37157, mean intensity of three independent experiments. **d**, Detection of H_2_O_2_ by live confocal microscopy in CytoHyPer-transfected BAECs either non-treated or treated with CGP in normoxia (Nx) or acute hypoxia (1% O_2_, Hp); representative images and time-course traces as mean of three independent experiments. **e**, Detection of ROS by live fluorescence microscopy with DCFDA in normoxia or transition from normoxia to acute hypoxia (2% O_2_); time-course traces and slopes, mean of three independent experiments. **f-g**, Superoxide detection by fluorescence microscopy with DHE in normoxia (Nx) or 10 min hypoxia (1% O_2_) or AA in primary **(f)** or immortalized **(g)** HUVECs; mean of three independent experiments. Two-tailed Student’s t-test for pairwise comparisons and one-way ANOVA and Tukey’s test for multiple comparisons n.s. not significant, * p<0.05, **<p0.01, *** p<0.001. **b-c**, statistical comparisons shown only for Nx vs 0-10 groups.

**Extended Data Fig. 4.**
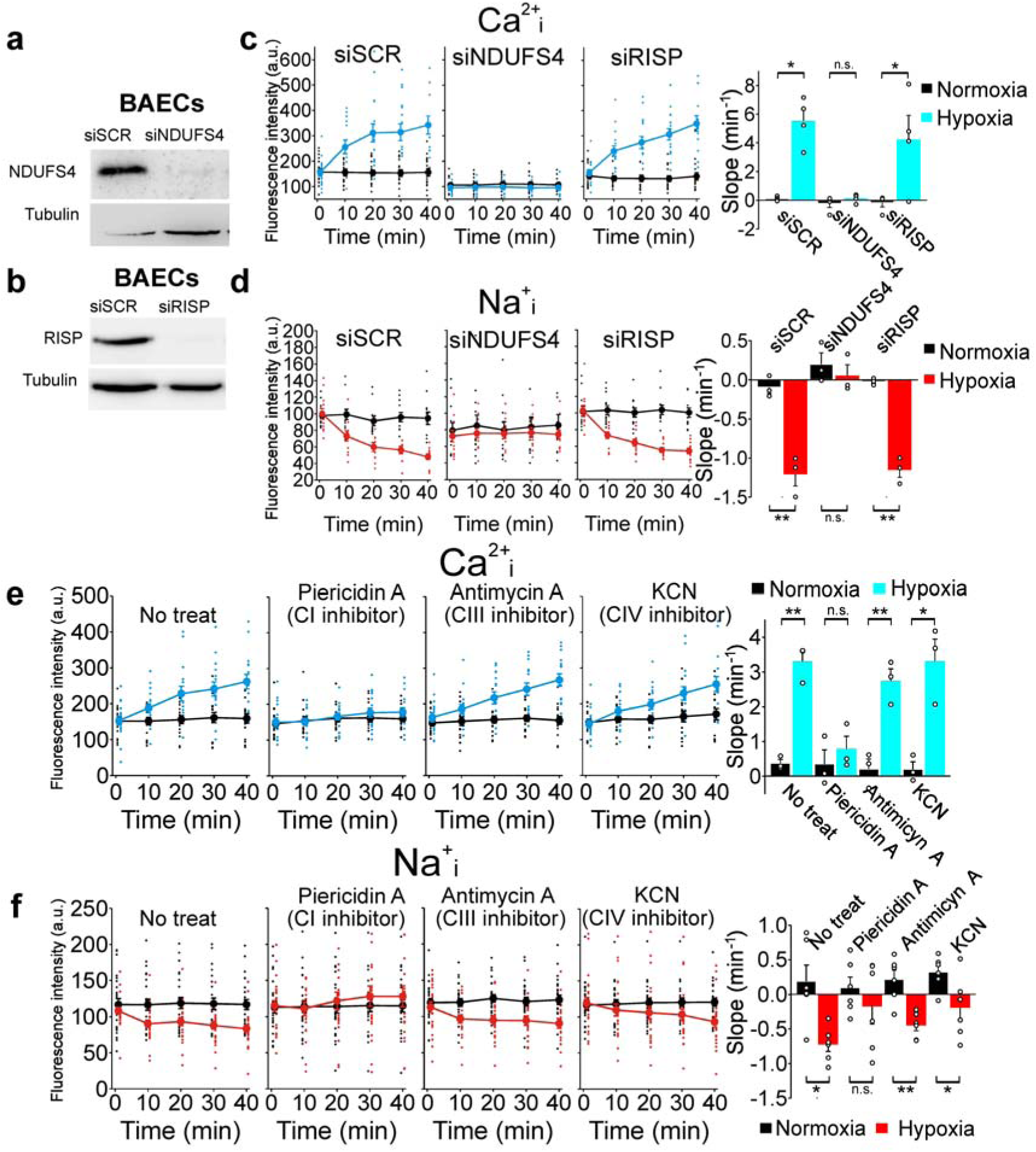
NCLX activation in acute hypoxia depends on mitochondrial complex I. **a, b**, Assessment of interference of subunits of CI (NDUFS4) or CIII (RISP) by Western blot in whole-cell extracts from BAECs. **c, d**, Cytosolic Ca^2+^ (Ca^2+^_i_) or Na^+^ (Na^+^_i_) were measured by live imaging confocal microscopy with Fluo-4 AM or CoroNa Green, respectively, in normoxia or acute hypoxia (1% O_2_), time-course traces and slopes, n=3 (Ca^2+^_i_), n=4 (Na^+^_i_). **e-f**, Effect of OXPHOS inhibitors on cytosolic Ca^2+^ (**e**, n=3) or Na^+^ (**f**, n=4) measured as in **c, d**. Two-tailed Student’s t-test. n.s. not significant, * p<0.05, **<p0.01.

**Extended Data Fig. 5.**
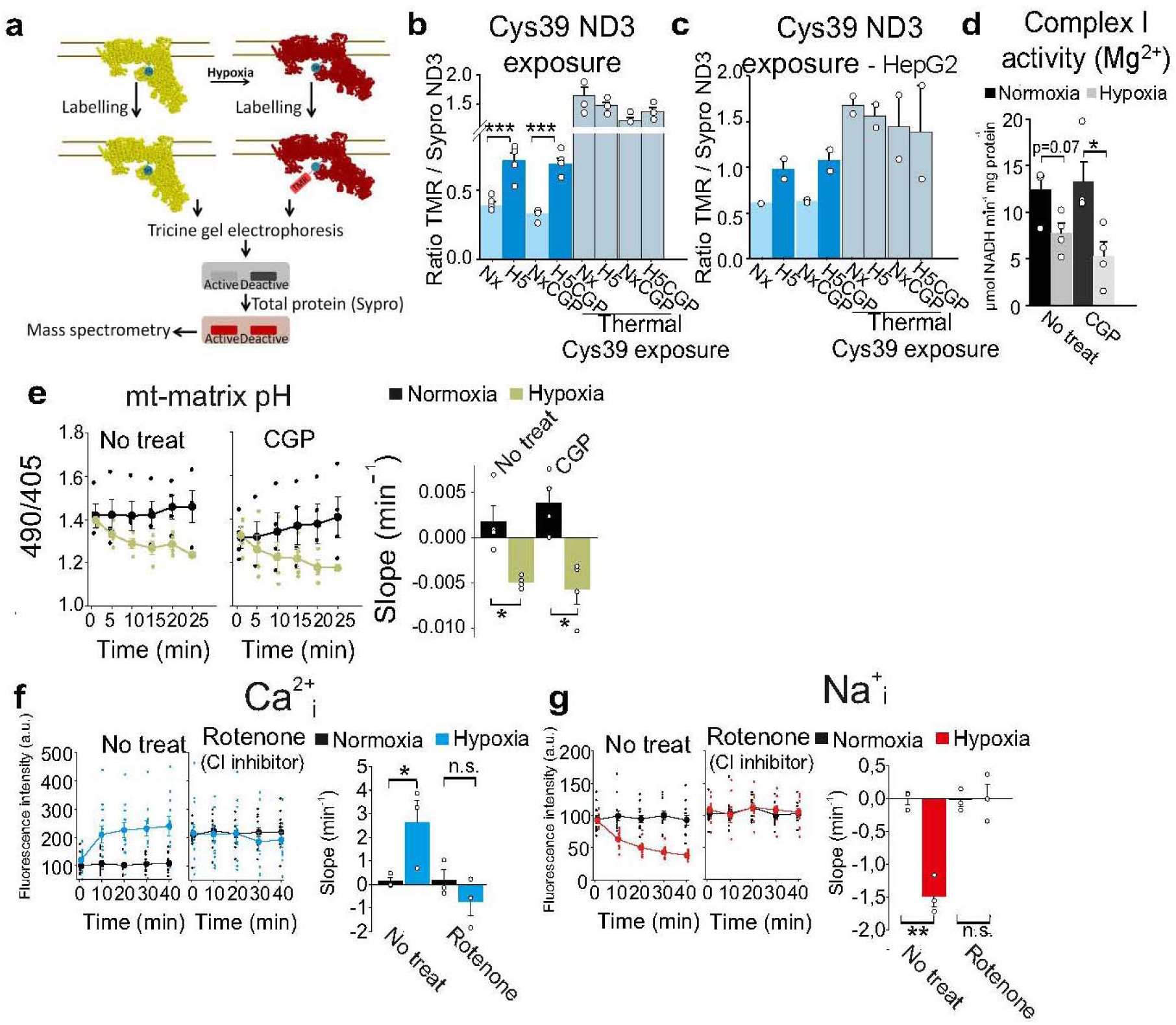
Acute hypoxia promotes A/D transition in complex I independently of NCLX activity. **a-c**, ND3•Cys39 exposure, which reflects the D state of CI, measured as the ratio between TMR signal (Cys39 labelling) and Sypro Ruby staining (total protein for the ND3 band, identified by mass spectrometry^11^). Thermal deactivation is used as a positive control of CI D state. **a**, scheme of the technique^11^; BAECs (**b**, n=4) or HepG2 (**c**, n=2) exposed to normoxia (Nx), 5 min of hypoxia (1% O_2_, H5), normoxia with CGP-37157 (NxCGP), 5 min of hypoxia with CGP-37157 (H5CGP). **d**, Complex I reactivation rate measured in the presence of Mg^2+^ in isolated mitochondrial membranes from BAECs subjected to normoxia, 10 min of hypoxia (1% O_2_) and NCLX inhibition with CGP-37157 (n=4). **e**, Mitochondrial matrix acidification induced by acute hypoxia (1% O_2_)^11^ and the effect of NCLX inhibition in BAECs transfected with mitosypHer, time-course traces and slopes (n=4). **f-g**, Effect of the CI inhibitor rotenone on cytosolic Ca^2+^ (**f**) or cytosolic Na^+^ (**g**) measured by live confocal microscopy with Fluo-4 AM or CoroNa Green, respectively, in BAECs in normoxia or acute hypoxia (1% O_2_), n=3. Two-tailed Student’s t-test for pairwise comparisons and one-way ANOVA with Tukey’s test for multiple comparisons. n.s. not significant, * p<0.05, **<p0.01, *** p<0.001.

**Extended Data Fig. 6.**
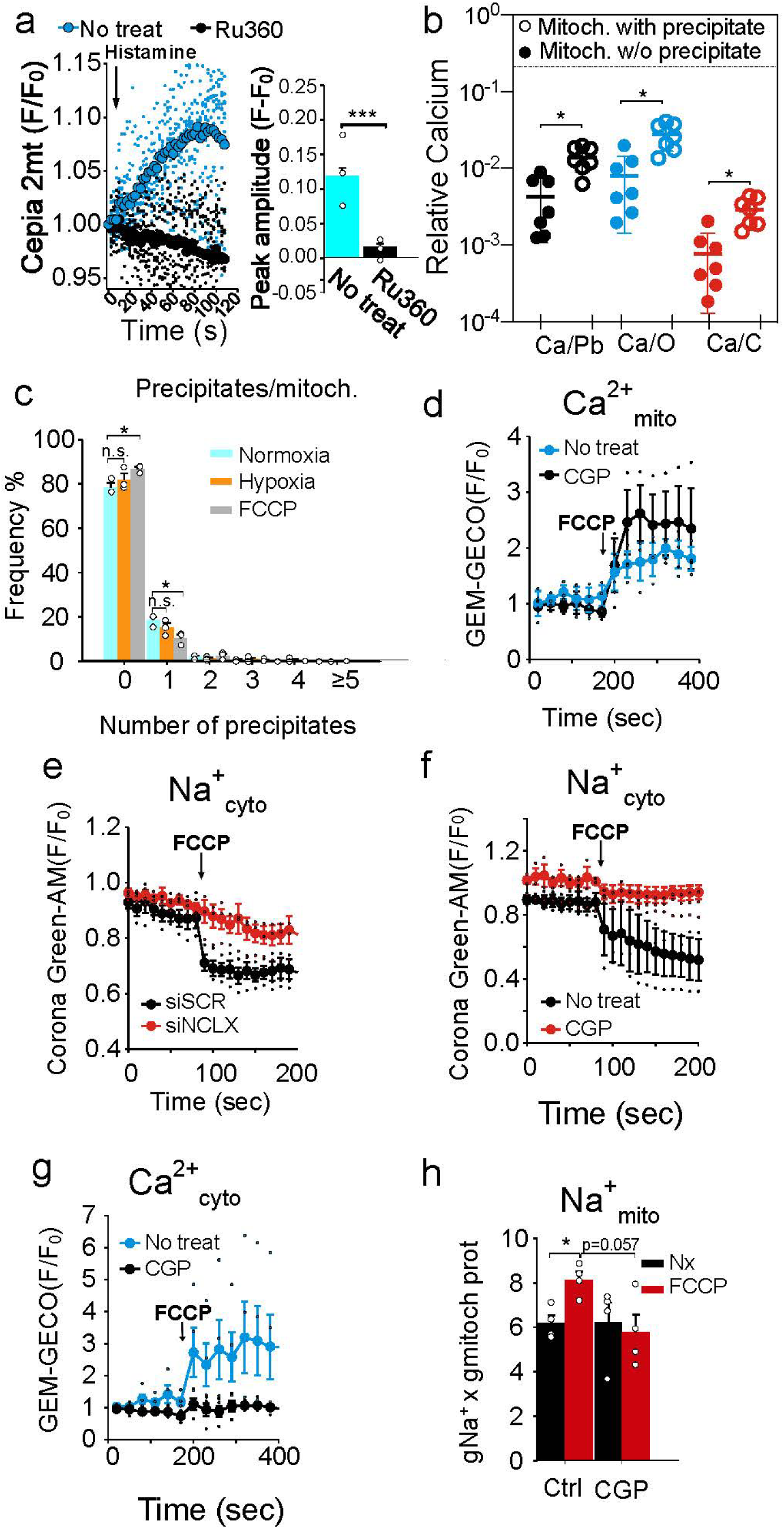
Mitochondrial matrix acidification promotes mitochondrial Na^+^/Ca^2+^ exchange via NCLX. **a**, Assessment of the effect of 1 µM Ru360 on mitochondrial Ca^2+^ influx rates in BAECs transfected with the mitochondria-directed Ca^2+^ reporter protein Cepia 2mt (n=3), after addition of histamine. Left, mean traces plotted as fluorescence change relative to initial fluorescence (F/F_0_); right, mitochondrial Ca^2+^ peak amplitude. **b**, TEM-EDX determination of calcium element (Ca) versus carbon (C), oxygen(O) and lead (Pb) content in regions of mitochondria with (empty circles) or without (filled circles) electron dense spots. **c**, Frequency of calcium phosphate precipitates per mitochondrion in BAECs, seen by transmission electron microscopy during normoxia (741 mitochondria), 10 min of hypoxia (1% O_2_; 619 mitochondria) or 30 min with 1 µM FCCP (393 mitochondria), sum of three independent experiments. **d**, Mitochondrial Ca^2+^ measured by live cell confocal microscopy in BAECs transfected with mito-GEM-GECO, either non-treated or treated with CGP-37157, before and after addition of 1 µM FCCP. **e, f**, Cytosolic Na^+^ measured by live cell confocal microscopy with CoroNa Green in BAECs transfected with siSCR or siNCLX **(e)**, non-treated BAECs or treated with CGP-37157 **(f)**, before and after addition of 1 µM FCCP. **f**, Cytosolic Ca^2+^ measured by live cell confocal microscopy in BAECs transfected with cyto-GEM-GECO, either non-treated or treated with CGP-37157, before and after addition of 1 µM FCCP. **h**, Mitochondrial Na^+^ measured by SBFI fluorimetry in mitochondria isolated from non-treated BAECs (Con) or treated with CGP-37157, FCCP or both. Fluorescence signal relative to starting signal (F/F0), n=3 (except **h**, n=4). One-way ANOVA with Tukey’s test. n.s. not significant, * p<0.05.

**Extended Data Fig. 7.**
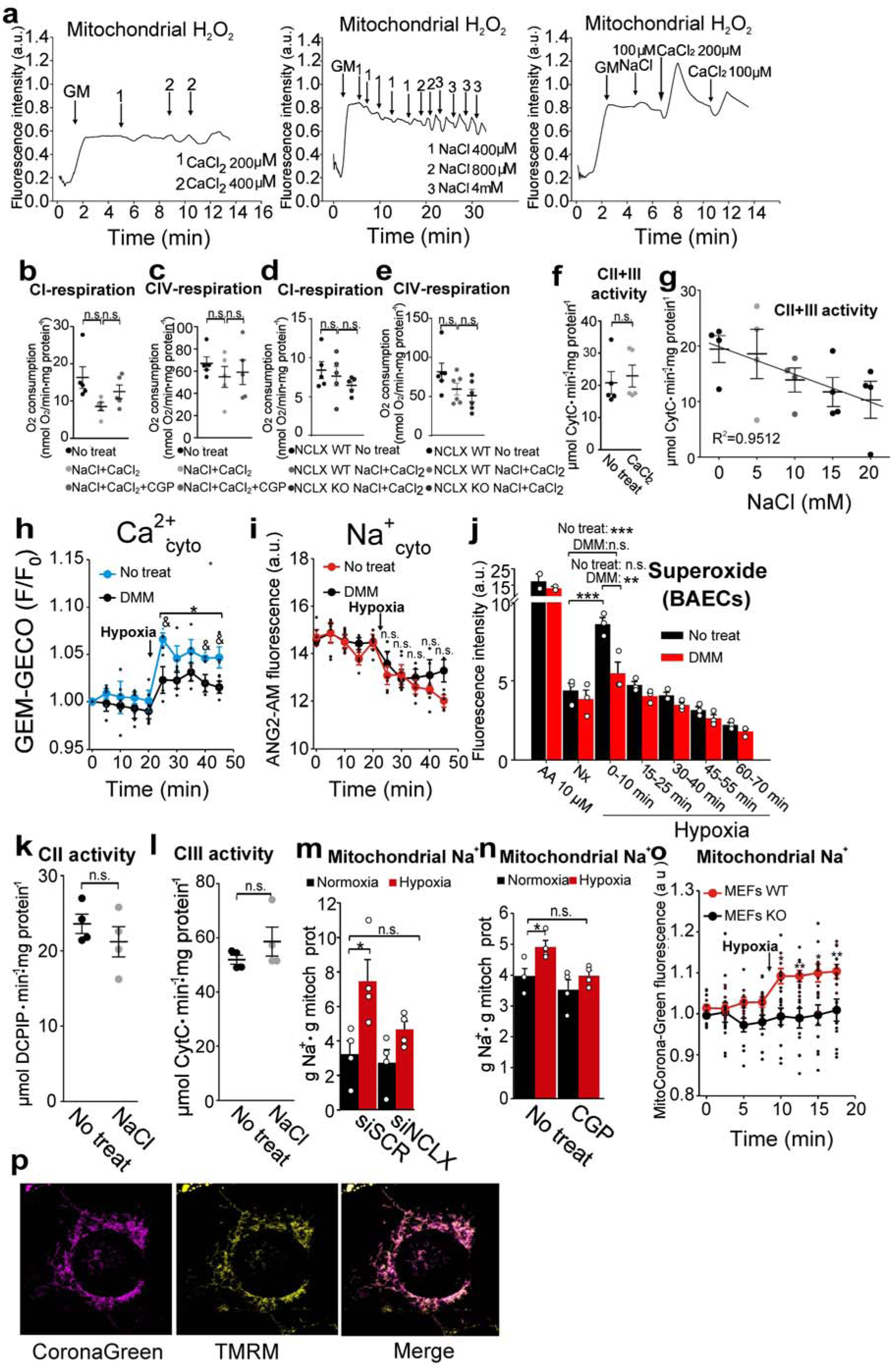
Mitochondrial Na^+^ import decreases OXPHOS and produces ROS. **a**, Effect of NaCl and/or CaCl_2_ additions on H_2_O_2_ production detected with Amplex Red in isolated rat heart mitochondria (500 µg) respiring after addition of glutamate/malate (GM) in KCl-EGTA buffer. Representative traces of five independent experiments. **b-e**, Effect of NCLX activation by 10 mM NaCl/0.1 mM CaCl_2_ on glutamate/malate- **(b, d)** or TMPD-based **(c, e)** OCR in isolated coupled mitochondria from BAECs **(b-c)** or MEFs **(d-e)** (n=5). **f, g**, Effect of 0.1 mM CaCl_2_ (n=5) or NaCl (n=4) additions on CII+III activity in isolated mitochondrial membranes from BAECs. **h**, Cytosolic Ca^2+^ measured by live cell confocal microscopy of BAECs transfected with cyto-GEM-GECO, either non-treated (No treat) or treated with 10 µM DMM during normoxia and hypoxia (1% O_2_; n=4). **i**, Cytosolic Na^+^ measured by live cell confocal microscopy with ANG2-AM in non-treated BAECs (No treat) or treated with 10 µM DMM during normoxia and hypoxia (1% O_2_; n=4). **j**, Superoxide detection by fluorescence microscopy after incubation with DHE in 10-min time windows in non-treated BAECs (No treat) or treated with 10 µM DMM during normoxia (Nx) or hypoxia (1% O_2_; n=3). **k-l**, Effect of 10 mM NaCl addition on succinate dehydrogenase activity (**k**) or ubiquinone 2-cytochrome *c* activity (**l**) from BAECs mitochondrial membranes (n=4). **m-n**, Effect of hypoxia (1% O_2_) in Na^+^_mito_ content in BAECs transfected with siSCR, siNCLX **(m)** or treated with CGP-37157 **(n)** (n=4). **o**, Effect of hypoxia (1% O_2_) in Na^+^_mito_ content in WT and KO MEFs, measured with CoroNa-Green adapted for mitochondrial loading and live fluorescence. **p**, Representative images showing colocalization of Corona Green-AM and TMRM signals after application of a long incubation protocol for Corona Green AM. Student’s t-test for pairwise comparisons and one-way ANOVA with Tukey’s test for multiple comparisons. n.s. not significant, * p<0.05, **<p0.01, *** p<0.001. Student’s t-test (No treat vs DMM): ^&^ p < 0.05. **j**, statistical comparisons shown only for Nx vs 0-10 groups. Pearson correlation coefficient in **g** (R=-0.9753).

**Extended Data Fig. 8.**
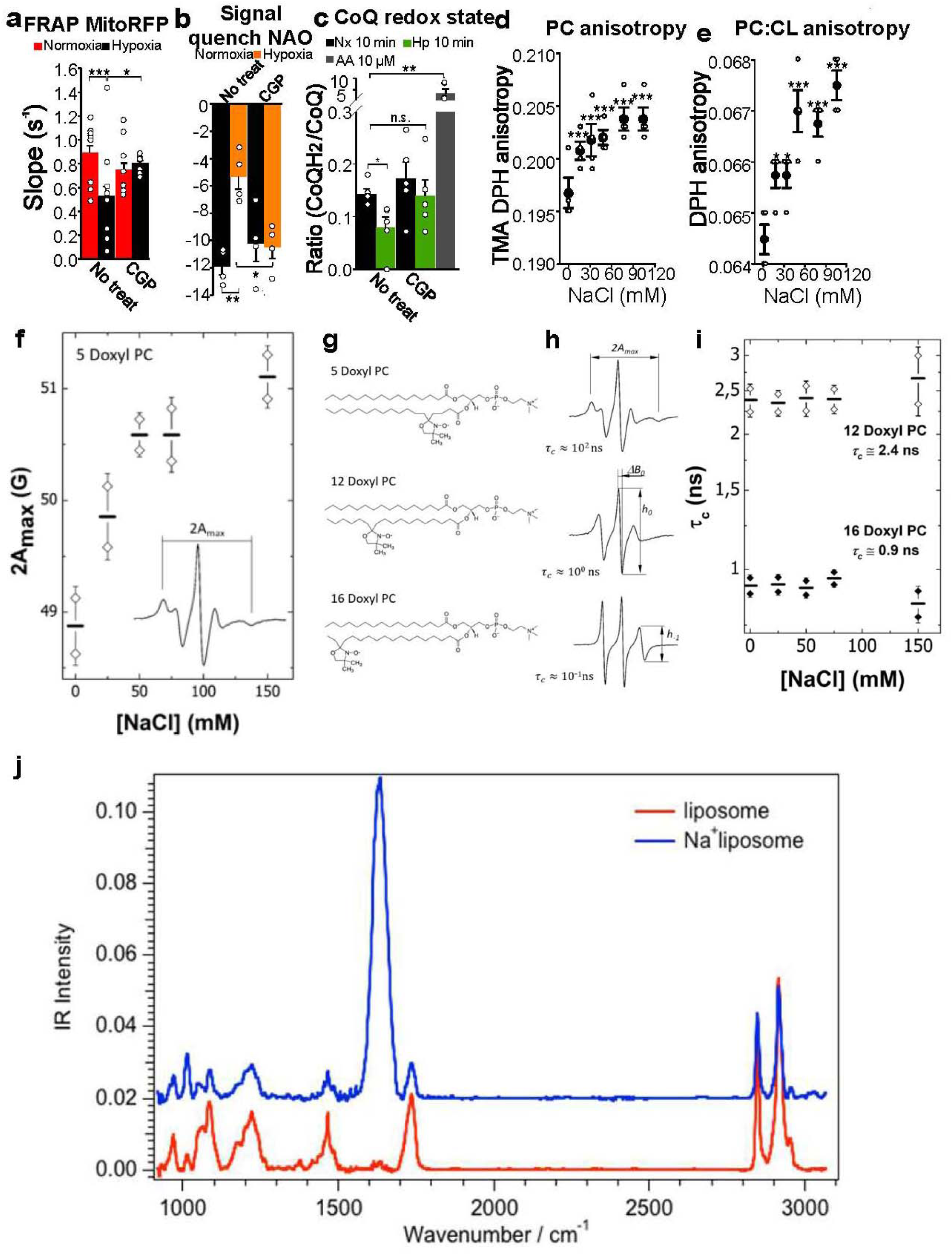
Mitochondrial Na^+^ import decreases inner mitochondrial membrane fluidity. **a**, FRAP of BAECs expressing mitoRFP in normoxia or hypoxia for 20 min (1% O_2_; n ≥ 6 determinations from four independent experiments), with or without CGP-37157. **b**, NAO quench signal of BAECs exposed to normoxia or hypoxia for 15 min (1% O_2_; n=5), with or without CGP-37157. **c**, CoQ_10_ redox state of HUVECs subjected to normoxia (Nx), 10 min of hypoxia (Hp 10 min; 1% O_2_), or treated with AA, n=4. **d-e**, Anisotropy of phosphatydilcholine (PC) or PC:cardiolipin (PC:CL) liposomes treated with increasing concentrations of NaCl measured by TMA-DPH or DPH fluorescence. **f-i**, ESR spectra of 5-, 12- and 16-Doxyl PC in DOPC liposomes. 5-Doxyl PC exhibited an increased restricted motion (broadening of the hyperfine splitting, 2A_max_) as a function of the NaCl concentration whereas the correlation time for 12- and 16-Doxyl PC remained unchanged. **f**, Chemical structures of 5-, 12- and 16-Doxyl PC. **g**, Hyperfine splitting (2A_max_) of 5-Doxyl PC measured by ESR in PC liposomes treated with increasing concentrations of NaCl (n=2). **h**, ESR of PC liposomes. **i**, rotational times, τ_c_, of 12- and 16-Doxyl PC in PC liposomes treated with increasing concentrations of NaCl as measured by ESR (n=2). **j**, IR spectroscopy absorption spectra of PC liposomes treated or not with 16 mM NaCl. Student’s t-test for pairwise comparisons and one-way ANOVA with Tukey’s test for multiple comparisons. n.s. not significant, * p<0.05, **<p0.01, *** p<0.001.

## References

1 Sena, L. A. & Chandel N. S. Physiological roles of mitochondrial reactive oxygen species. Mol Cell 48, 158–167, (2012).

2 Shadel, G. S. & Horvath T. L. Mitochondrial ROS signaling in organismal homeostasis. Cell 163, 560–569, (2015).

3 Dan Dunn, J., Alvarez, L. A., Zhang, X. & Soldati T. Reactive oxygen species and mitochondria: A nexus of cellular homeostasis. Redox Biol 6, 472–485, (2015).

4 Chouchani, E. T. et al. Ischaemic accumulation of succinate controls reperfusion injury through mitochondrial ROS. Nature 515, 431–435, (2014).

5 Guzy, R. D. & Schumacker P. T. Oxygen sensing by mitochondria at complex III: the paradox of increased reactive oxygen species during hypoxia. Exp Physiol 91, 807–819, (2006).

6 Fernández-Agüera, M. C. et al. Oxygen Sensing by Arterial Chemoreceptors Depends on Mitochondrial Complex I Signaling. Cell Metab 22, 825–837, (2015).

7 Hernansanz-Agustín, P. et al. Acute hypoxia produces a superoxide burst in cells. Free Radic Biol Med 71, 146–156, (2014).

8 Sylvester, J. T., Shimoda, L. A., Aaronson, P. I. & Ward J. P. Hypoxic pulmonary vasoconstriction. Physiol Rev 92, 367–520, (2012).

9 Giorgi, C., Marchi, S. & Pinton P. The machineries, regulation and cellular functions of mitochondrial calcium. Nat Rev Mol Cell Biol 19, 713–730, (2018).

10 Bers, D. M., Barry, W. H. & Despa S. Intracellular Na+ regulation in cardiac myocytes. Cardiovasc Res 57, 897–912, (2003).

11 Hernansanz-Agustín, P. et al. Mitochondrial complex I deactivation is related to superoxide production in acute hypoxia. Redox Biol 12, 1040–1051, (2017).

12 Luongo, T. S. et al. The mitochondrial Na+/Ca2+ exchanger is essential for Ca2+ homeostasis and viability. Nature 545, 93–97, (2017).

13 Hamanaka, R. B. & Chandel N. S. Mitochondrial reactive oxygen species regulate cellular signaling and dictate biological outcomes. Trends Biochem Sci 35, 505–513, (2010).

14 Babot, M., Birch, A., Labarbuta, P. & Galkin A. Characterisation of the active/de-active transition of mitochondrial complex I. Biochim Biophys Acta 1837, 1083–1092, (2014).

15 Zhu, J., Vinothkumar, K. R. & Hirst J. Structure of mammalian respiratory complex I. Nature 536, 354–358, (2016).

16 Fiedorczuk, K. et al. Atomic structure of the entire mammalian mitochondrial complex I. Nature 538, 406–410, (2016).

17 Greenawalt, J. W., Rossi, C. S. & Lehninger A. L. Effect of Active Accumulation of Calcium and Phosphate Ions on the Structure of Rat Liver Mitochondria. J Cell Biol 23, 21–38, (1964).

18 Wolf, S. G. et al. 3D visualization of mitochondrial solid-phase calcium stores in whole cells. Elife 6, (2017).

19 Chandel, N. S. et al. Reactive oxygen species generated at mitochondrial complex III stabilize hypoxia-inducible factor-1alpha during hypoxia: a mechanism of O2 sensing. J Biol Chem 275, 25130–25138, (2000).

20 Guzy, R. D. et al. Mitochondrial complex III is required for hypoxia-induced ROS production and cellular oxygen sensing. Cell Metab 1, 401–408, (2005).

21 Carafoli, E., Tiozzo, R., Lugli, G., Crovetti, F. & Kratzing C. The release of calcium from heart mitochondria by sodium. J Mol Cell Cardiol 6, 361–371, (1974).

22 Cox, D. A. & Matlib, M. A. A role for the mitochondrial Na(+)-Ca2+ exchanger in the regulation of oxidative phosphorylation in isolated heart mitochondria. J Biol Chem 268, 938–947, (1993).

23 Acín-Pérez, R., Fernández-Silva, P., Peleato, M. L., Pérez-Martos, A. & Enríquez, J. A. Respiratory active mitochondrial supercomplexes. Mol Cell 32, 529–539, (2008).

24 Lenaz, G. & Genova M. L. Mobility and function of coenzyme Q (ubiquinone) in the mitochondrial respiratory chain. Biochim Biophys Acta 1787, 563–573, (2009).

25 Budin, I. et al. Viscous control of cellular respiration by membrane lipid composition. Science 362, 1186–1189, (2018).

26 Petit, J. M., Maftah, A., Ratinaud, M. H. & Julien, R. 10N-nonyl acridine orange interacts with cardiolipin and allows the quantification of this phospholipid in isolated mitochondria. Eur J Biochem 209, 267–273, (1992).

27 Gallet, P. F., Petit, J. M., Maftah, A., Zachowski, A. & Julien R. Asymmetrical distribution of cardiolipin in yeast inner mitochondrial membrane triggered by carbon catabolite repression. Biochem J 324 (Pt 2), 627–634, (1997).

28 Horvath, S. E. & Daum G. Lipids of mitochondria. Prog Lipid Res 52, 590–614, (2013).

29 Sarewicz, M. & Osyczka A. Electronic connection between the quinone and cytochrome C redox pools and its role in regulation of mitochondrial electron transport and redox signaling. Physiol Rev 95, 219–243, (2015).

30 Pabst, G. et al. Rigidification of neutral lipid bilayers in the presence of salts. Biophys J 93, 2688–2696, (2007).

31 Böckmann, R. A., Hac, A., Heimburg, T. & Grubmüller, H. Effect of sodium chloride on a lipid bilayer. Biophys J 85, 1647–1655, (2003).

32 Cordomí, A., Edholm, O. & Perez J. J. Effect of ions on a dipalmitoyl phosphatidylcholine bilayer. a molecular dynamics simulation study. J Phys Chem B 112, 1397–1408, (2008).

33 Michelakis, E. D., Thebaud, B., Weir, E. K. & Archer S. L. Hypoxic pulmonary vasoconstriction: redox regulation of O2-sensitive K+ channels by a mitochondrial O2-sensor in resistance artery smooth muscle cells. J Mol Cell Cardiol 37, 1119–1136, (2004).

34 Moreno, L. et al. Ceramide mediates acute oxygen sensing in vascular tissues. Antioxid Redox Signal 20, 1–14, (2014).

35 Desireddi, J. R., Farrow, K. N., Marks, J. D., Waypa, G. B. & Schumacker P. T. Hypoxia increases ROS signaling and cytosolic Ca(2+) in pulmonary artery smooth muscle cells of mouse lungs slices. Antioxid Redox Signal 12, 595–602, (2010).

36 Connolly, M. J., Prieto-Lloret, J., Becker, S., Ward, J. P. & Aaronson P. I. Hypoxic pulmonary vasoconstriction in the absence of pretone: essential role for intracellular Ca2+ release. J Physiol 591, 4473–4498, (2013).

37 Lapuente-Brun, E. et al. Supercomplex assembly determines electron flux in the mitochondrial electron transport chain. Science 340, 1567–1570, (2013).

38 Enríquez, J. A. Supramolecular Organization of Respiratory Complexes. Annu Rev Physiol 78, 533–561, (2016).

39 Letts, J. A., Fiedorczuk, K., Degliesposti, G., Skehel, M. & Sazanov L. A. Structures of Respiratory Supercomplex I+III2 Reveal Functional and Conformational Crosstalk. Mol Cell, (2019).

40 Navarro-Antolín, J., Rey-Campos, J. & Lamas S. Transcriptional induction of endothelial nitric oxide gene by cyclosporine A. A role for activator protein-1. J Biol Chem 275, 3075–3080, (2000).

41 Muñoz, C. et al. Transcriptional up-regulation of intracellular adhesion molecule-1 in human endothelial cells by the antioxidant pyrrolidine dithiocarbamate involves the activation of activating protein-1. J Immunol 157, 3587–3597, (1996).

42 McCombs, J. E. & Palmer A. E. Measuring calcium dynamics in living cells with genetically encodable calcium indicators. Methods 46, 152–159, (2008).

43 Cogliati, S. et al. Mechanism of super-assembly of respiratory complexes III and IV. Nature 539, 579–582, (2016).

44 Schägger, H. Tricine-SDS-PAGE. Nat Protoc 1, 16–22, (2006).

45 Zhang, H. Thin-Film Hydration Followed by Extrusion Method for Liposome Preparation. Methods Mol Biol 1522, 17–22, (2017).

46 Rouser, G., Fkeischer, S. & Yamamoto A. Two dimensional then layer chromatographic separation of polar lipids and determination of phospholipids by phosphorus analysis of spots. Lipids 5, 494–496, (1970).

47 Scorrano, L. et al. A distinct pathway remodels mitochondrial cristae and mobilizes cytochrome c during apoptosis. Dev Cell 2, 55–67, (2002).

48 Scialo, F. et al. Mitochondrial ROS Produced via Reverse Electron Transport Extend Animal Lifespan. Cell Metab 23, 725–734, (2016).

49 Rodríguez-Aguilera, J. C., Cortés, A. B., Fernández-Ayala, D. J. & Navas P. Biochemical Assessment of Coenzyme Q10 Deficiency. J Clin Med 6, (2017).

50 Yubero, D. et al. Secondary coenzyme Q10 deficiencies in oxidative phosphorylation (OXPHOS) and non-OXPHOS disorders. Mitochondrion 30, 51–58, (2016).

51 Stone, T. J., Buckman, T., Nordio, P. L. & McConnell, H. M. Spin-labeled biomolecules. Proc Natl A cad Sci U S A 54, 1010–1017, (1965).

52 Martínez-Ruiz, A. et al. RNase U2 and alpha-sarcin: a study of relationships. Methods Enzymol 341, 335–351, (2001).

53 Gasset, M., Martínez del Pozo, A., Oñaderra, M. & Gavilanes J. G. Study of the interaction between the antitumour protein alpha-sarcin and phospholipid vesicles. Biochem J 258, 569–575, (1989).

