## Supplementary videos legends for "Mitochondrial Na^+^ controls oxidative phosphorylation and hypoxic redox signalling"

**Supplementary video 1. FRAP of normoxic WT MEFs expressing mito-RFP.** Time-lapse showing a representative FRAP of wild type MEFs expressing mito-RFP subjected to normoxia.

**Supplementary video 2. FRAP of hypoxic WT MEFs expressing mito-RFP.** Time-lapse showing a representative FRAP of wild type MEFs expressing mito-RFP subjected to 10 min of hypoxia (1% O_2_).

**Supplementary video 3. FRAP of normoxic KO MEFs expressing mito-RFP.** Time-lapse showing a representative FRAP of NCLX KO MEFs expressing mito-RFP subjected to normoxia.

**Supplementary video 4. FRAP of hypoxic KO MEFs expressing mito-RFP.** Time-lapse showing a representative FRAP of NCLX KO MEFs expressing mito-RFP subjected to hypoxia (1% O_2_).

**Supplementary video 5. FRAP of normoxic KO+pNCLX MEFs expressing mito-RFP.** Time-lapse showing a representative FRAP of NCLX KO MEFs expressing pNCLX and mito-RFP, subjected to normoxia.

**Supplementary video 6. FRAP of normoxic KO+pNCLX MEFs expressing mito-RFP.** Time-lapse showing a representative FRAP of NCLX KO MEFs expressing pNCLX and mito-RFP, subjected to hypoxia (1% O_2_).

**Supplementary video 7. FRAP of normoxic WT+dnNCLX MEFs expressing mito-RFP.** Time-lapse showing a representative FRAP of wild type MEFs expressing dnNCLX and mito-RFP, subjected to normoxia.

**Supplementary video 8. FRAP of hypoxic WT+dnNCLX MEFs expressing mito-RFP.** Time-lapse showing a representative FRAP of wild type MEFs expressing dnNCLX and mito-RFP, subjected to 10 min of hypoxia (1% O_2_).
